# Sex significantly impacts the function of major depression-linked variants *in vivo*

**DOI:** 10.1101/2021.11.01.466849

**Authors:** Bernard Mulvey, Din Selmanovic, Joseph D. Dougherty

## Abstract

Genome-wide association studies have discovered blocks of common variants—likely transcriptional-regulatory—associated with major depressive disorder (MDD), though the functional subset and their biological impacts remain unknown. Likewise, why depression occurs in females more frequently than males is unclear. We therefore tested the hypothesis that risk-associated functional variants interact with sex and produce greater impact in female brains. We developed methods to directly measure regulatory variant activity and sex interactions using massively parallel reporter assays (MPRAs) in the mouse brain *in vivo*, in a cell type-specific manner. We measured activity of >1,000 variants from >30 MDD loci, identifying extensive sex-by-allele effects in mature hippocampal neurons and suggesting sex-differentiated impacts of genetic risk may underlie sex bias in disease. Unbiased informatics approaches indicated that functional MDD variants recurrently disrupt sex hormone receptor binding sequences. We confirmed this with MPRAs in neonatal brains, comparing brains undergoing the masculinizing hormone surge to hormonally-quiescent juveniles. Our study provides novel insights into the influence of age, biological sex, and cell type on regulatory-variant function, and provides a framework for *in vivo* parallel assays to functionally define interactions between organismal variables like sex and regulatory variation.

**One-Sentence Summary:** Massively parallel assays *in vivo* identified extensive functional and sex-interacting common variants in depression risk loci.

## Introduction

Major depressive disorder (MDD) is a profoundly disruptive and sometimes lethal disorder, affecting women 2-3 times more frequently than men across countries and cultures (*1*). Sex differences are present across multiple levels of the disease, from symptom profiles (*2*) and effective drug classes (*3*) to brain-wide gene expression (*4*, *5*). Genome-wide association studies(GWASes) have identified dozens of linkage regions each containing numerous single-nucleotide polymorphisms (SNPs) associated with MDD, demonstrating its heritability (*6*–*8*). More recently, sex-by-genotype (SxG) analyses of large GWAS cohorts have revealed that MDD risk loci are for men and women, yet these loci explain up to 4-fold greater MDD heritability in females (*9*, *10*). These findings suggest that sex interacts with a common pool of SNPs to attenuate or amplify the MDD risk they confer. However, disease-associated SNPs are seldom found in protein-coding space, complicating prediction of their molecular consequences. Instead, these SNPs are found in probable regulatory elements (REs), including transcriptional-regulatory sequences predicted from measures such as chromatin marks, accessibility, and conformation. Specific brain regions and cell types are enriched for such measures at—and in putative regulatory target genes of—MDD-associated loci, including the hippocampus (*7*, *11*) and excitatory neurons (*12*–*16*), suggesting sites of action for these REs. In particular, there has been long-standing interest in the hippocampus regarding both MDD pathology and sex differences in the brain. Hippocampal volume reductions in MDD patients have been widely reported (*17*). Moreover, the hippocampus is subject to influences of sex from perinatal (*18*, *19*) to adult life, presenting in MDD as sex differences in hippocampal volume loss (*20*) and gene expression (*4*).

Determining the identity of functional SNPs from MDD-associated regions is the first key step toward understanding the biological perturbations resulting from risk genotypes, which can in turn enable inference of dysregulated target genes and shared regulatory programs involved across loci. However, studies connecting MDD-associated SNPs to gene expression in brain tissue, even those that considered sex effects (*21*), have been limited to indirect and imperfect indicators of function (e.g., chromatin state), or confounded by linkage disequilibrium (e.g., for expression quantitative trait loci (eQTLs)). In contrast, *direct measurement* of regulatory output of common variants associated with disease has largely been restricted to the *in vitro* setting. The large-scale *in vitro* identification of functional regulatory variants has been made possible by massively parallel reporter assays (MPRAs), a method for functionally detecting activity from thousands of REs (and their variants) simultaneously. In brief, MPRAs adapt a traditional reporter assay paradigm—placing REs upstream of an optically measured reporter (e.g., luciferase)—but adds a unique, RE-identifying “barcode” sequence to the reporter’s 3’ untranslated region, enabling quantification of activity for thousands of REs simultaneously by RNA barcode sequencing. MPRAs have enabled identification of trait- and disease-associated SNPs affecting REs in culturable, disease-relevant cell types *in vitro* (*22*–*25*). However, the complexities of cell types interacting in the brain and of the sex hormonal milieu cannot readily be emulated *ex vivo*.

We overcome these prior limitations in functional regulatory SNP (rSNP) identification to interrogate the biological contexts under which MDD rSNPs act by delivering an MPRA library of MDD-associated variants (*26*) into the adult mouse hippocampus *in vivo*. Building on prior brain MPRAs of enhancers (*27*) and an RE variant (*28*), our approach greatly extends *in vivo* MPRA methods to identify rSNPs and their sex interactions, including those which are cell type-specific. First, we combined MPRA with translating ribosome affinity purification (TRAP) to simultaneously identify MDD rSNPs in both excitatory neurons and the broader hippocampus; these experiments utilized mice of both sexes, enabling us to additionally test the hypothesis that rSNPs are subject to sex-by-genotype (SxG) interactions. Finally, to further characterize the potential role of circulating hormones in sex-differentiated rSNP activity, and to functionally replicate predicted fetal brain RE enrichments suggesting a role for MDD SNPs during circuit organization (*15*, *29*–*32*), we likewise delivered the library to the mouse brain *in utero.* This allowed us to identify rSNPs neonatally, coinciding with a testosterone surge and critical period for establishing sex-specific brain circuitry (*33*), and test for loss of SxG effects in juveniles, when hormonal influences are quiescent. In sum, we illustrate that MPRAs can be leveraged *in vivo* to directly identify not only functional variants, but their context-dependence on age, sex, and cell type, while demonstrating that all three of these factors have substantial impacts on MDD-associated regulatory variation.

## Results

### Combining Translating Ribosome Affinity Purification (TRAP) with MPRAs

Adeno-associated viruses (AAVs) have long been used to evaluate the activity of single REs in the brain, most recently in MPRA-like designs to screen multiple REs in parallel with RNA-seq-based quantification (*27*, *34*, *35*). This suggests it may also be possible to adapt AAV-MPRAs to study functional consequences of RE variants associated with disease. As MDD genetic risk is enriched in neuronal REs, we tested the feasibility of combining a cell type-specific profiling method, TRAP, with MPRA to attain measurements of RE activity specifically in neurons. We generated 4 small AAV9 libraries (**Fig. 1A-C)** expressing dsRed under the control of the *hsp68* minimal promoter, with full-length human promoters (denoted *pGene*) with documented expression in neurons (human *pCAMK2A*), astrocytes (*pGFAP*), or all cells (*pPGK2*), each carrying unique 3’ untranslated region (UTR) barcodes for quantification by RNA-seq. We first individually confirmed cell-type specificity by immunofluorescence (IF) (**Fig. 1D-F**), and then delivered (*36*) a mixed pool of the four barcoded AAV9 libraries into the brain of postnatal day 2 (P2) neuron-specific TRAP mice (*Snap25-Rpl10a-eGFP)* (*37*). TRAP was then used to compare total brain and neuronal activity levels of the three promoters and the *hsp68* promoter alone. We prepared MPRA sequencing libraries from 1) the delivered AAV pool (DNA), 2) total brain (input) RNA, and 3) TRAP (neuronal) RNA and assessed expression of each barcode by calculating a ratio of RNA counts to DNA counts. Results were highly replicable at the level of barcodes (**Fig. 1G-H**) in both RNA fractions. Moreover, neuronal TRAP fractions demonstrated increased expression of barcodes driven by *pCAMK2A* and lower expression of *pGFAP*-paired barcodes (**Fig. 1I**). These results demonstrated that cell type-specific effects, even of relatively small magnitudes, can be detected using a combined MPRA-TRAP approach. We then turned to applying this method to test the effects of human variants associated with psychiatric disease.

**Figure 1.**
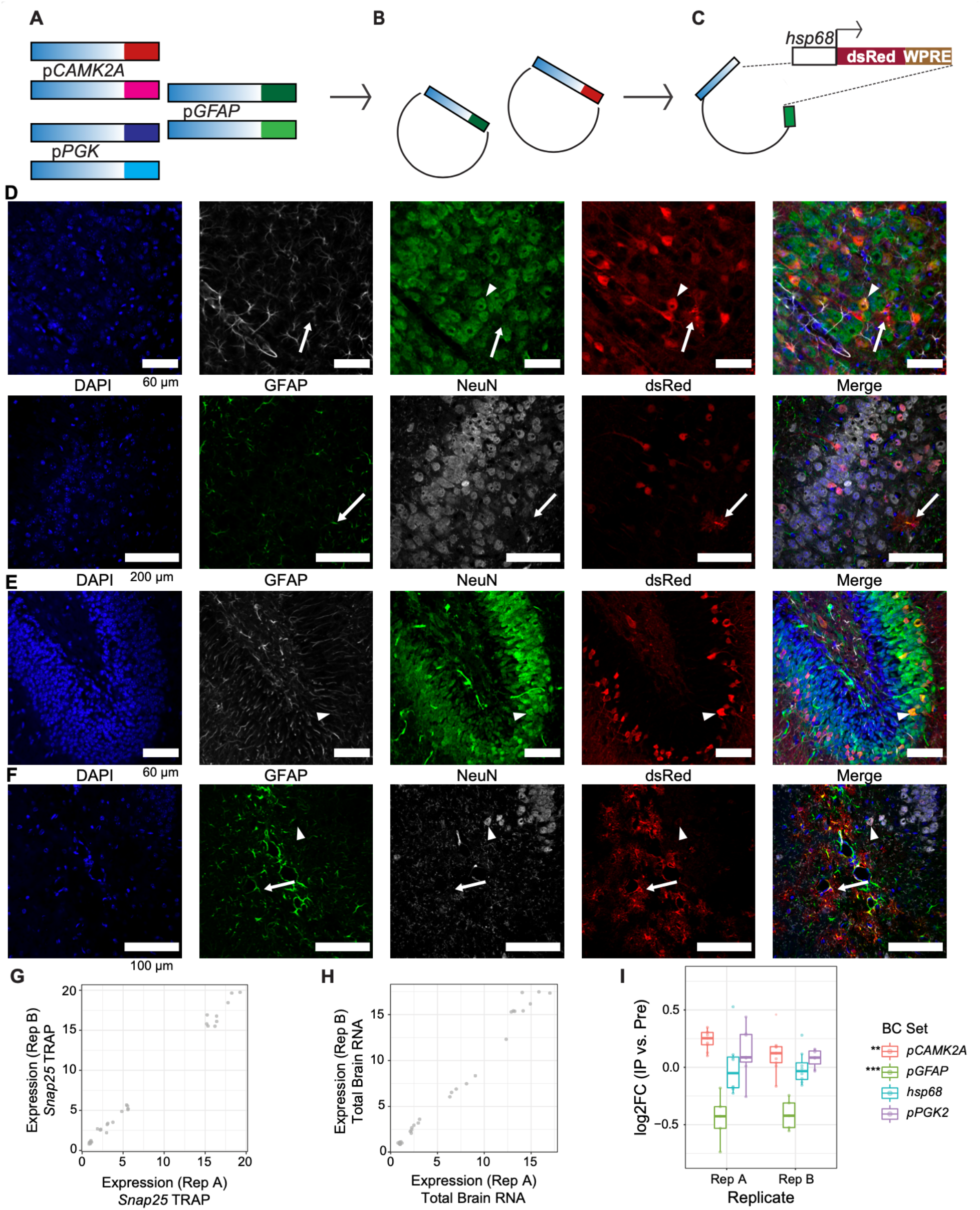
Proof-of-principle: cell type-specific MPRA *in vivo*. (**A**) Cell type-specific promoters, *p*GFAP (astrocytic) and *p*CAMK2A (neuronal) were barcoded by PCR. (**B**) Amplicons were cloned into an AAV plasmid. **C)** Further restriction cloning added a reporter cassette containing a minimal *hsp68* promoter, dsRed, and an RNA-stabilizing 3’ UTR hepatitis E “woodchuck” (WPRE) element. Barcoded pools with each promoter were packaged into AAV9 separately. (**D-F**) IF of P27 mouse brain after P2 injection with a single AAV9 barcode pool. (D) *hsp68* promoter alone preferentially drove dsRed expression in neurons, while (E) *p*CAMK2A drove reporter expression solely in neurons, and (F) *p*GFAP drove predominantly astrocytic dsRed expression. (**G**) Replicability of barcode expression for the SNAP25-TRAP RNA fraction (Pearson’s *r*=0.9975) and (**H**) total brain tissue (input) (*r*=0.9927). (**I**) TRAP expression, compared to total brain expression, was higher for *pCAMK2A* and lower for *pGFAP*, as expected. ^**^*p* ≤ 5 × 10^−4^, ^***^*p* ≤ 5 × 10^−8^

### Identification of rSNPs and their sex-allele interactions in total hippocampus and excitatory neurons

Given the association of MDD with sex, hippocampal pathology, and neuronal genetics, we sought to identify regulatory SNPs among variants selected from broad linkage disequilibrium (LD) regions associated with MDD in total hippocampus. To further investigate the roles of SNPs and sex in hippocampal excitatory neurons in particular, we performed these experiments in a cross of Slc17a7 (or, Vglut1)-*Cre* mice (*38*) to a *Cre* recombinase-dependent TRAP mouse line (*37*).

The analyzed MPRA library covered 40 GWAS loci spanning ~1,000 SNPs in LD R^2^>0.1 with MDD-associated tag variants. SNPs were prioritized by their overlap with human brain and neural cell type eQTLs, histone marks, enhancer RNA overlap, and chromatin contacts (see *Data Availability*). Of these SNPs, 926 were from 29 MDD GWAS loci (*6*, *7*, *32*, *39*–*43*), 19 were from 2 loci identified by meta-analysis of MDD and autism spectrum disorders (*44*), and 21 were from 4 loci for MDD-correlated traits (mood instability, anxiety, and neuroticism) (*45*–*48*). 126bp human genomic sequences centered on the SNP were generated for each allele and paired to 10 unique 10bp barcodes per allelic sequence for internal replication (**Fig. S1**). These were inserted into an AAV plasmid, then cloned to contain an *hsp68* minimal promoter (*49*) driving dsRed along with the “woodchuck” hepatitis 3’UTR element to improve recovery of reporter RNAs (*50*), and packaged into AAV9.

We delivered the AAV9 library bilaterally into the hippocampus of Vglut1-TRAP mice ages P60-P80 (n=6 per sex), followed by hippocampal dissection and TRAP (**Fig. 2A**) to identify rSNPs and their shared regulatory features (**Fig. 2B**). IF of hippocampi from additional mice confirmed robust hippocampal expression of the dsRed reporter 28 days after injection **(Fig. 2C**). We first confirmed by qPCR that TRAP RNA was depleted for glial marker genes several-fold as expected (**Fig. 2D**), then conducted MPRA sequencing (**Table S1)** and analysis of both input RNA (hippocampus) and TRAP RNA (Vglut1^+^) for each sample.

**Figure 2.**
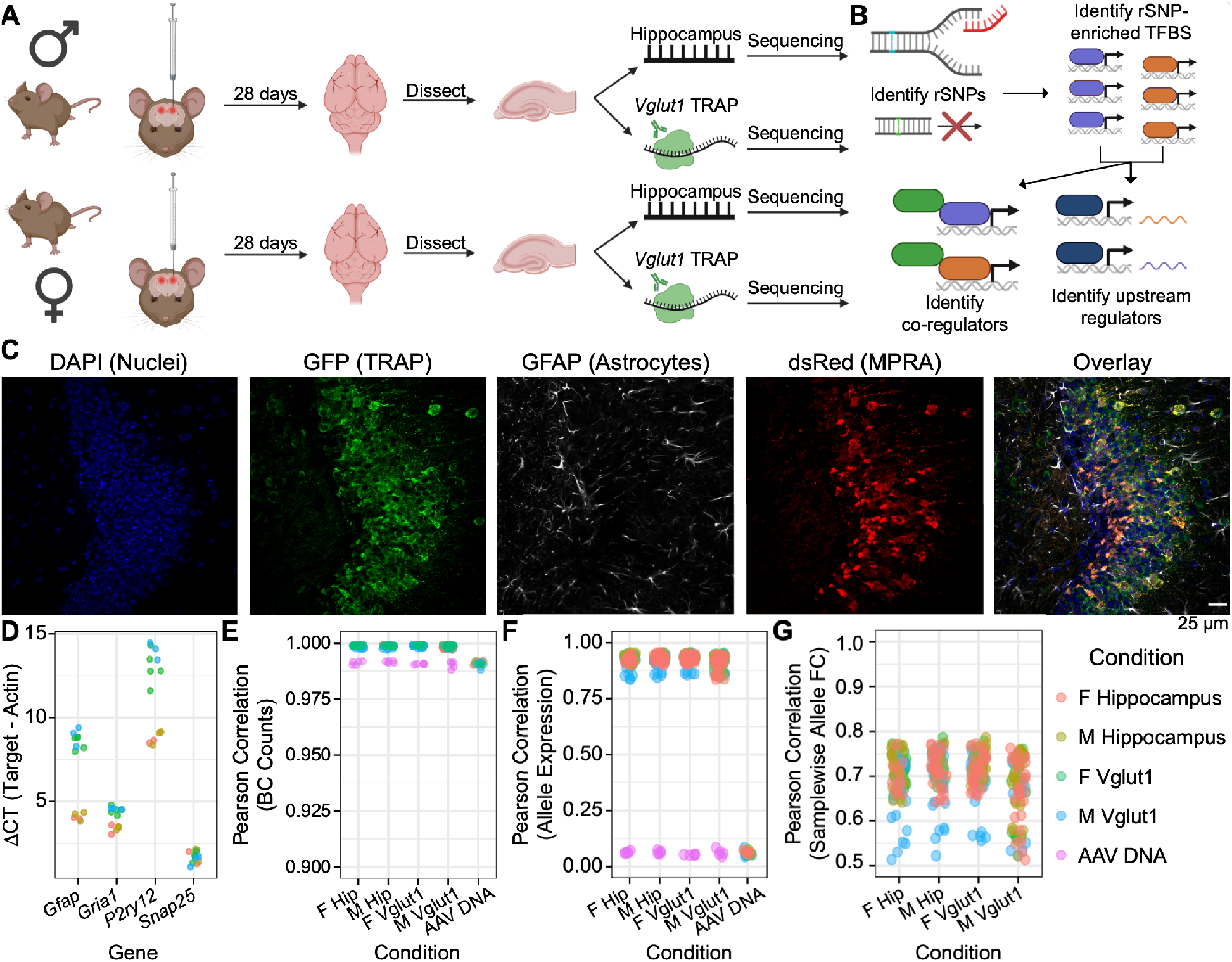
Experimental design, analysis plan, and quality control: adult mouse hippocampus and its excitatory neurons. (**A**) Adult male and female C57BL/6J mice received bilateral stereotactic injections into the hippocampus delivering the AAV9-packaged MPRA. TRAP yielded two RNA fractions per sample: “input” (total hippocampal) and TRAP (Vglut1^+^). (**B**) Analyses identified regulatory SNPs (rSNPs), transcription factor (TF) binding sites (TFBSes) enriched at rSNPs, shared protein interactors among these TFs, and shared regulators of these TFs’ expression. (**C**) IF of Vglut1-TRAP mouse hippocampus 28 days after MPRA-AAV9 delivery, illustrating strong TRAP (GFP) co-expression with dsRed reporter, confirming RNA from the latter is present in the cell type of interest. (**D**) qPCR confirmed depletion of glial genes (*Gfap, P2ry12*) and modest enrichment of excitatory neuron marker *Gria1* in Vglut1^+^ RNA. (**E**) Barcode count correlations between replicates. Each point represents the cross-correlation between one sample of the type on the x-axis and one of the color-coded type. (**F**) Correlation of mean barcode expression between replicates. (**G**) Correlation of samplewise allelic differences in expression. PCC: Pearson correlation coefficient (or Pearson’s *r*).

After quality control, n=5 male and n=5 female hippocampal samples (both the input and TRAP RNAs) were retained for MPRA analysis covering ~1,000 SNPs (see *Online Methods*). These showed replicability at the level of barcode counts (pairwise Pearson’s *r* 0.7-0.89) (**Fig. 2E**), RE-level expression (*r* 0.85-0.96) (**Fig. 2F**), and allelic fold changes within each condition (*r* 0.90-0.94) (**Fig. 2G**), confirming we were able to reliably measure SNP-mediated regulatory effects and differences from a defined cell type *in vivo*.

#### rSNPs in the adult hippocampus and hippocampal Vglut1^+^ neurons

We first assessed ~1,000 SNPs for allelic effects in each individual sex and RNA fraction, using linear mixed models (LMMs) fitting barcode expression as a function of allele with random barcode effects (see *Online Methods* and **Fig. S2-S4**). We calculated empirical *p* (*p*_emp_) values using 50,000 simulated ‘allelic’ comparisons between subsets of random barcodes coupled to the minimal promoter alone (*26*, *51*) to account for technical and barcode-mediated noise. Significance was called at FDR-corrected *p*_emp_<0.2, a stringency comparable to a recent study of sex-interacting eQTLs (*21*). **Supplemental Data S1** provides allelic beta (log2FC) values and significance status for variants at 10 different significance thresholds.

In total hippocampus, we identified 36 (male) and 31 (female) rSNPs, 34 and 31 of which were from MDD loci, respectively. While male and female total hippocampi had similar numbers of rSNPs, we observed a striking sex difference in the number of rSNPs in Vglut1^+^ cells specifically—only 7 (male) compared to 58 (female), indicating that within excitatory neurons, a higher proportion of MDD SNPs have discernible allelic effects in females.. Moreover, all 7 male rSNPs were also functional in females. Notable rSNPs from ≥1 condition included rs2563323 and rs250427, putative brain and hippocampal (*52*) eQTLs for *SRA1*, a noncoding RNA which activates nuclear receptors even in the absence of their ligand (*53*). Also notable were rs301806—an MDD GWAS index SNP (*40*)—and rs301807 (**Fig. 3A-B**), both of which likely regulate the nearby gene*RERE* (*15*, *54*).

**Figure 3.**
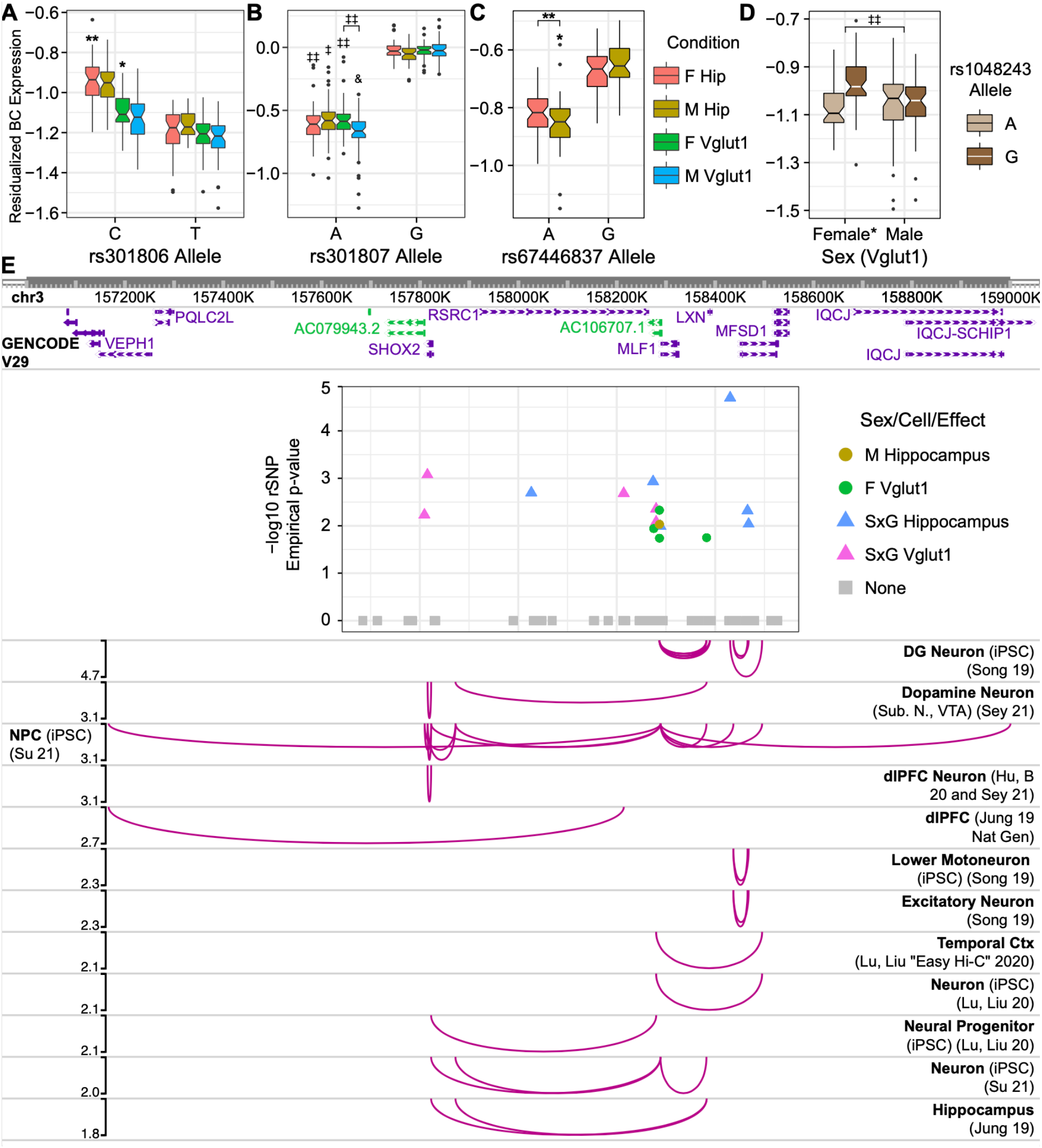
Adult hippocampus rSNPs and complex context-dependent, polygenic architecture of the *RSRC1* locus. Boxplots show single-barcode (BC) expression levels adjusted for random effects across analyzed replicates. Center bars: median; boxes: 25-75% quantile; whiskers: observations spanning box edges to ± 1.5*interquantile range (IQR); single points: observed values outside whisker range. Notched regions span ± 1.58 * IQR / sqrt (n measurements), approximating the 95% confidence interval for comparing median BC expression (*55*). rSNPs corresponding to *RERE* locus (*32*) tag variants (**A**) rs301806 and (**B**) rs301807 are shown. (**C, D**) Sex-interacting SNPs from the *RSRC1* locus (*39*) included (C) rs67446837 and (D) rs1048243. (**E**) rSNPs identified in the *RSRC1* locus and their regulatory target genes in human tissues ascertained by Hi-C. A plot of rSNP effects, colored by most significant condition, is embedded with its x-axis in human genomic (hg19) coordinates; chromatin contacts between the SNPs and distal gene promoters are illustrated below (*56*–*61*). Curve heights correspond to −log10*p*_emp_ for the plotted rSNP. iPSC: Induced pluripotent stem cell; DG: dentate gyrus; Ctx: cortex. *: *p_emp_*-derived FDR < 0.25; **: <0.2; ‡ < 0.15; ‡‡ < 0.1; & < 0.05.

#### Sex-interacting rSNPs in hippocampus

Given the role of sex in MDD risk and the observed differences in rSNP activity in the adult hippocampus, we sought to investigate whether sex interacts with MDD risk genotypes. To ensure there were not confounding sex differences in minimal promoter activity, we compared minimal promoter-only barcode expression between sexes for the two tissue fractions, finding no sex difference in activity (*t*-test of barcode expression, p>0.5 in both comparisons; **Fig. S5**).

We therefore performed combined-sex LMM analysis of the Vglut1^+^ and hippocampus results, identifying 41 sex-allele interaction rSNPs in each. Notably, while only 1 SxG rSNP was shared between total hippocampus and Vglut1^+^, sex-interacting SNPs originated from the same GWAS loci across both analyses. The tag locus rs1193510 (*39*), for example, contained 11 sex-interacting rSNPs, including several unique to hippocampus (**Fig. 3C**) or Vglut1^*+*^ (**Fig. 3D**). This region is rich in human neural chromatin contacts (*56*–*61*), implicating several target genes of the identified rSNPs. Interestingly, SxG effects in hippocampus and Vglut1^+^ segregated into distinct portions of this LD region (**Fig. 3E**). We additionally examined three SNPs in our assay that were recently reported significant in sex-genotype interaction GWASes of over 500 traits (*62*). One of these reported variants, rs2400075, was found to have several significant sex-interacting associations to body traits; this variant is located near *LIN28B*—which shows sex-differential expression in mouse norepinephrine neurons (*63*)–and was a near-significant SxG rSNP in Vglut1^*+*^ (0.20<FDR<0.25).

#### Transcriptional-regulatory systems shared across hippocampal rSNPs

We next asked whether there were any shared transcriptional-regulatory mechanisms underlying MDD rSNP effects in the hippocampus. We tested whether rSNPs perturbed specific transcription factor (TF) binding motifs more frequently than expected by chance (defined by their rates in rSNPs vs. non-effect SNPs in the assay) (*26*). We assessed motif disruptions using motifbreakR (*64*) and RSAT var-tools (*65*) (see *Online Methods*) defining rSNPs at a nominal LMM *p*_emp_ of 0.05. This resulted in sets of 80-110 MPRA-identified rSNPs per condition (*p*_emp_ FDR levels of 0.22-0.28; **Data S2A**), ensuring adequate depth for enrichment analysis—akin to other methods which loosen significance thresholds to analyze higher-order relationships of granular molecular measures, e.g., polygenic risk scoring or gene ontology. To refine these results, we filtered the enriched TFs (FDR<0.05) to those with altered putative binding sites ≥4 rSNPs and expressed in Genotype-Tissue Expression atlas (v8) hippocampus in the corresponding sex.

Altogether, we identified 34 enriched TFs in male total hippocampus rSNPs and 19 in female. The hippocampal TFs identified were largely distinctive between sexes; for example, KLF family TFs were unique to male hippocampal rSNPs, while nuclear receptor (NR) TFs were mostly unique to female rSNPs (**Fig. 4A**). Among Vglut1^+^ rSNPs, we identified 8 TFs in female and 16 in male, many of which were shared (e.g., DLX1, POU3F1/2/3). POU3F2 has been previously shown to be a highly centralized, cross-disorder hub gene in postmortem brain co-expression analysis by PsychENCODE (*66*).

**Figure 4.**
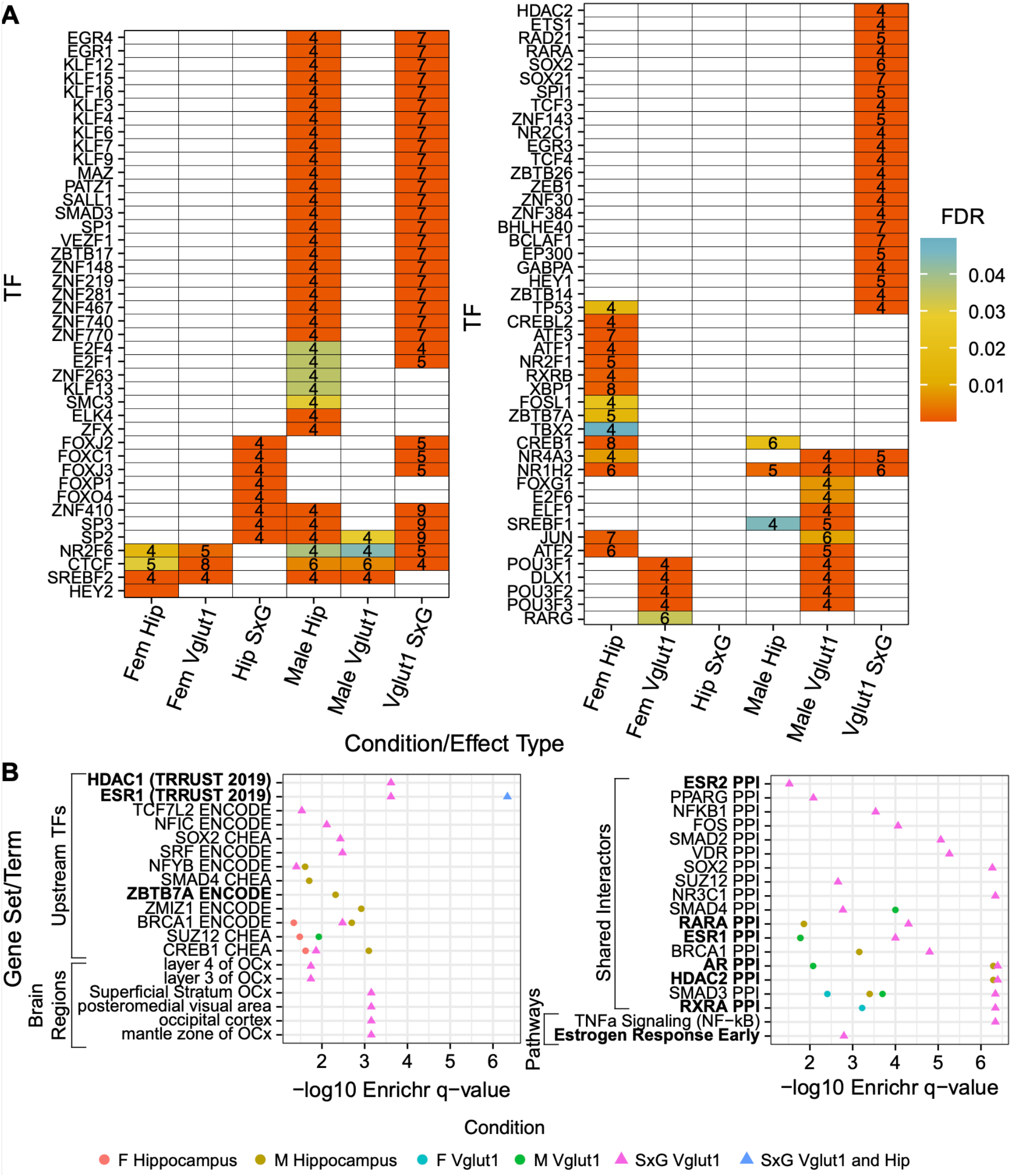
Shared regulatory architecture of rSNPs by sex, cell type, and sex-interacting SNP type. (**A**) TFs with binding motif disruptions by ≥ 4 nominally significant (empirical *p* < 0.05) rSNPs or sex-interacting rSNPs, enriched relative to nonfunctional SNPs. Number of rSNPs associated to a given binding site are shown. (**B**) Terms from Enrichr (*67*) analysis, identifying shared upstream regulators (TFs controlling expression of several of the TFs in panel A), brain regions enriched for expression of the rSNP-enriched TFs, protein interactors enriched among rSNP-enriched TFs, and MSigDB pathway term enrichment for rSNP-enriched TF sets. Bolded enrichments are discussed in the text.

To understand integrative biological functions of these TF sets, we utilized the tool Enrichr (*67*) for each TF set, including both tissues per sex, to identify putative *upstream* regulators, co-interacting TFs (protein-protein interactions (PPIs)), enriched disease gene sets, ontologies, and brain regions expressing the TFs (**Data S2B-C**). The most striking enrichments (> 25% of query TFs) for male glutamatergic and hippocampal TFs combined were for regulators *of* these TFs’ expression, including *CREB1* (14/48), *BRCA1* (19/48), and *ZBTB7A* (12/48). Male hippocampal TFs were likewise enriched for several upstream regulators, including *TCF3* (8/38), *HDAC2* (8/38), and *ZMIZ1* (9/38). *ZMIZ1* has roles in coactivation of androgen receptor (*AR*) (*68*) as well as *SMAD3* (*69*), consistent with enrichment of these TFs in *SMAD3* (6/38) and *AR* (7/38) PPIs. TFs from male glutamatergic neurons were enriched for four PPIs: *SMAD3* and *SMAD4* (both 5/16), consistent with MPRA signal from neuronal enrichment by TRAP, and more surprisingly, sex hormone receptors: estrogen receptor α (*ESR1*; 4/16) and *AR* (3/16).

Likewise, female glutamatergic TFs were enriched for PPIs with *SMAD3* (3/8), along with *RXRA* (3/8), replicating *in vivo* our recent *in vitro* findings of retinoid-interacting rSNPs in MDD-associated loci (*26*). Female total hippocampal TFs, on the other hand, were enriched for upstream regulation by several TFs, including *ATF2* (9/19), *BRCA1* (8/19), and *TCF3* (5/19).

#### Transcriptional-regulatory systems implicated in SxG interactions at MDD rSNPs in hippocampus

Using a similar approach to that above, we investigated TFs enriched at sex-allele interaction rSNPs (*p*_emp_<0.05, corresponding to FDRs of 0.34-0.37) relative to sex-agnostic rSNPs (combined-sexes LMM allele main effect *p*_emp_<0.05), and separately enriched to nonfunctional SNPs, then combined the two enrichment outputs to form TF sets (see *Online Methods*) (**Fig. 4A**). We identified only 8 TFs enriched among total hippocampal SxG rSNPs, including *ZNF410* and five TFs with a shared binding motif for *FOX*(*C1*/*J2/J3/O4/P1*). However, we identified 54 TFs enriched at glutamatergic SxG rSNPs, including six of those from total hippocampus, suggesting coherent regulatory dynamics spanning MDD loci in this cell type. SxG TFs included several *KLF* members, as well as nuclear receptors (*NR4A3, NR2C1, NR1H2*), and another retinoid TF, *RARA*.

We ran both tissue fraction SxG TF sets through Enrichr, though nothing notable appeared for total hippocampus given the small size of the set. Top glutamatergic SxG TF enrichments included upstream regulation by *BRCA1 (*21/57) and *CREB1* (15/57), as well as PPIs with *AR* (11/57), *ESR1* (11/57), and *HDAC2* (12/57) (**Fig. 4B**). Also enriched was an MsigDB functional gene set term, “estrogen response early” (5/57). These SxG regulators implicate sex hormones and histone acetylation in both establishing sex-divergent MDD risk from upstream, e.g. via *ESR1* (5/57), and actuating it downstream via rSNP-enriched TFs and their protein interactors (*AR*, *ESR1*, *HDAC2*, *ZMIZ1*, *ZBTB7A*). Incidentally, both sex hormones (*70*) and histone (de-)acetylases (*71*) have been major areas highlighted in recent reviews of sex differences in MDD and mouse models thereof.

### Identification of rSNPs in developing whole mouse brain

As sex differences in brain structure and transcriptional regulation are established in part by the effects of sex hormones, including the perinatal testosterone surge (*33*), we sought to investigate whether MDD risk variants were subject to sex-differential regulation during early development. To be able to assess the brain during this period, we delivered the AAV library intracerebroventricularly to embryonic day 15 (E15) mice, followed by whole brain collection at postnatal day 0 (P0) and P10 (see *Online Methods*). P0—while not amenable to regionally-targeted viral assays—is in the midst of the testosterone surge, during which both acute activational effects and transcriptional-regulatory organizational effects occur; by contrast, P10 is a timepoint where sex hormones are effectively absent in normal development.

We first verified by IF that dsRed expression was detectable at P0 and P10 following *in utero* delivery. Clear, widespread reporter expression was apparent at both timepoints despite the relatively short incubation time (**Fig. 5A-D**; **Fig. S6**). IF at P10 demonstrated prominent expression of the reporter in the hippocampus (**Fig. 5D**)—a structure neither present at E15 nor well-developed at P0—consistent with prior observations of AAV9 expression ultimately occurring in hippocampus when delivered to the perinatal brain (*73*). We subsequently collected the whole brain (except cerebellum) for RNA isolation and MPRA sequencing. Additionally, we isolated DNA from n=4 brain samples (one per age and sex) to profile the transduced library contents, verifying that the distribution of delivered MPRA barcodes was similar both between replicates (*r* 0.86-0.94) and to input virus (*r* 0.88-0.91) (**Fig. E**). Ultimately, we analyzed 15 samples for P0 (6 female, 9 male) and 13 for P10 (6 female, 7 male; see *Supplemental Text*). Replicates from each condition had well-correlated barcode counts (PCC 0.86-0.98) and RE expression values (PCC 0.58-0.96) (**Fig. 5F-H**).

**Figure 5.**
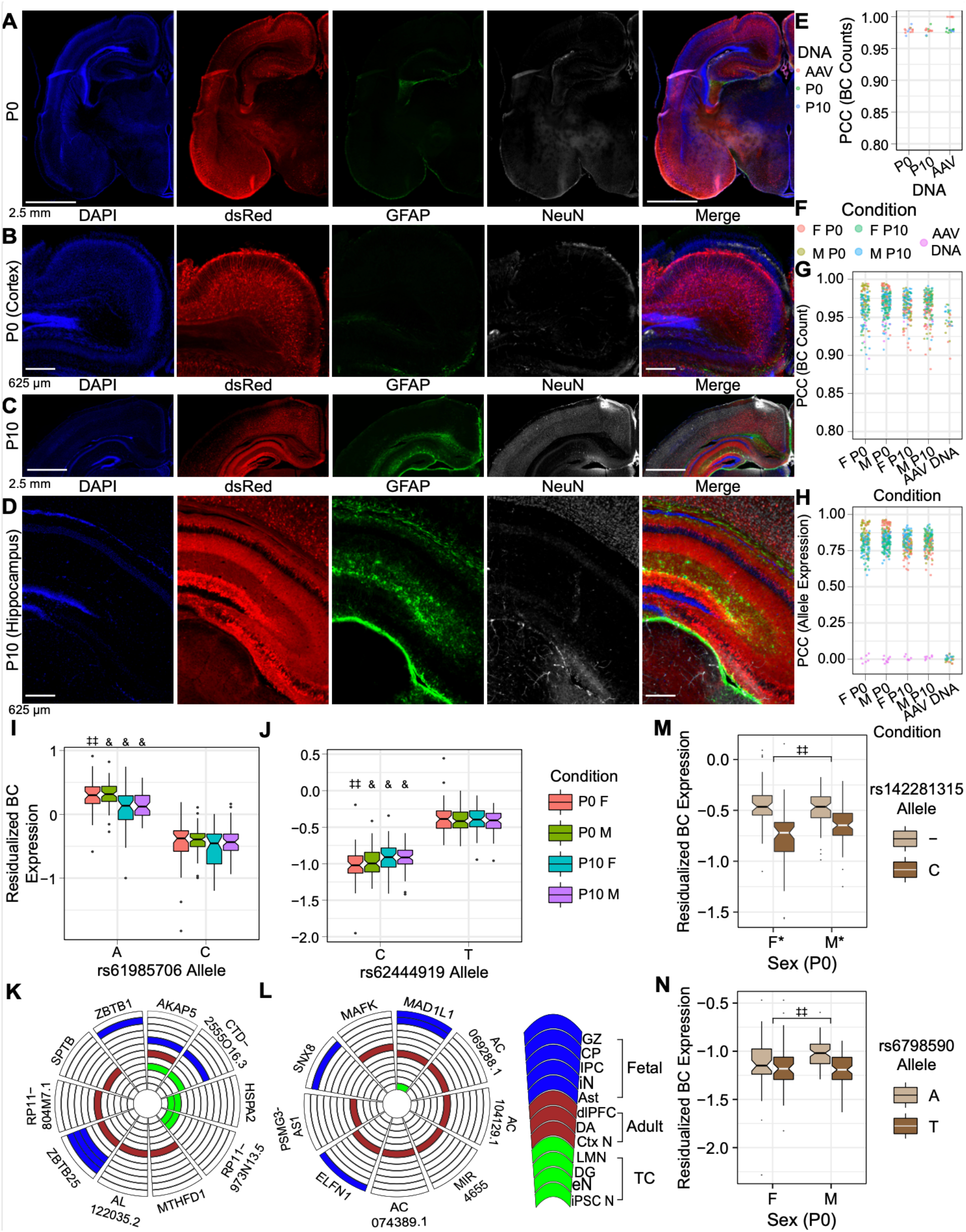
Validating the *in utero* MPRA delivery method, and identification of rSNPs and sex-interacting rSNPs in the developing brain. (**A, B**) IF of P0 brain after E15 MPRA-AAV delivery. (**C, D**) IF of P10 brain after MPRA-AAV delivery. (**E**) Comparability of barcode counts in recovered brain DNA and original AAV. (**F**) Color legend for panels G-H. (**G**) BC count correlation between samples. (**H**) Sequence expression correlation between samples. (**I, J**) rSNPs (I) rs61985706 and (J) rs62444919 (**J**) showed effects consistent across sexes and ages. (**K, L**) Putative target genes of the respective rSNPs from Hi-C in human fetal, adult, and cultured neural tissues. (**M**) Example P0 SxG SNP with comparatively small sex difference in allele effect size. (**N**) Example P0 SxG SNP with magnitude of sex difference in allelic effect comparable to smaller (female) allelic effect itself. GZ: germinal zone; CP: cortical plate (*72*); IPC: intermediate progenitor cell; iN: inhibitory neuron (*15*); Ast: astrocyte (*58*); dlPFC: dorsolateral prefrontal cortex; DA: dopamine neurons of substantia nigra and ventral tegmental area (*56*); Ctx N: cortical neuron (*61*); LMN: lower motor neuron; eN: excitatory neuron (*58*); iPSC N: iPSC-derived neuron (*59*). *: *p*_emp_-derived FDR < 0.25; **: <0.2; ‡ < 0.15; ‡‡ < 0.1; & < 0.05.

Within single sexes at P0, we identified 5 rSNPs in females and 12 rSNPs in males, respectively (*p*_emp_ FDR < 0.2), 4 of which were shared between conditions with consistent effect direction. By contrast, we identified 105 female and 72 male rSNPs at P10, with 42 rSNPs identified in both conditions with consistent direction of effect. 4 of these shared rSNPs were the same rSNPs shared between P0 sexes, with consistent effect direction in all four conditions (two of which are illustrated, **Fig. 5I, J**). Three of these four rSNPs also have rich chromatin contact evidence supporting gene regulatory roles in fetal, adult, and cultured human neural cell types (for the illustrated rSNPs, **Fig. 5K** and **5L, respectively**), consistent with their detection as rSNPs in whole brain tissue, highlighting *in utero* MPRA delivery as a robust method for detecting functional variation in the developing brain.

#### Sex-allele interactions are widespread neonatally but absent during hormonal quiescence

To identify SxG interactions occurring during neurodevelopment, we tested for SxG interactions as before, now within age groups. Again, the minimal promoter-only control was not sex-differentially expressed (*t*-test of barcode expression, *p*>0.1) (**Fig. S7**). At *p*_emp_ FDR <0.2, we identified 31 rSNPs with sex interactions in the P0 brain (e.g., **Fig. 5M-N)**. By contrast, we identified *no* SxG interactions at *p*_emp_ FDR<0.25 among the 930 analyzed SNPs at P10. We confirmed this result—despite similar *n* and inter-sample correlations to P0—by repeating the SxG analysis with the least variable 5 samples per sex (removing two males and one female) (**Data S3**).

#### Transcriptional-regulatory systems implicated in brain-wide rSNP function in postnatal development

We examined single-sex, single-age rSNP sets for enrichment of TF motif perturbations using motifbreakR and RSAT approaches as before, and again using nominally significant (*p*_emp_<0.05) rSNPs as the set tested for enrichment. We did not filter enriched TFs for expression in analogous human tissue, as fetal whole-brain gene expression profiles are unavailable; we instead required a more stringent minimum of 5 rSNPs to be found at significantly enriched (FDR<0.05) TF motifs.

In P0 brain, we identified 20 TFs enriched at female rSNPs and 43 at male rSNPs, 9 of which were shared (**Fig. 6A**). Shared transcription factors included *EGR1/4*, *TBX1/15*, and *IRF9*. Male-specific transcription factors again included *ZBTB7A* and *NR4A3* (**Fig. 6A**). Female-specific TFs included a variety of zinc finger TFs, Krüpell-like factors (*KLF*s), endothelial-developmental regulator *SOX17*, and the neurodevelopmental TF *SMAD3* (**Fig. 6A**).

**Figure 6.**
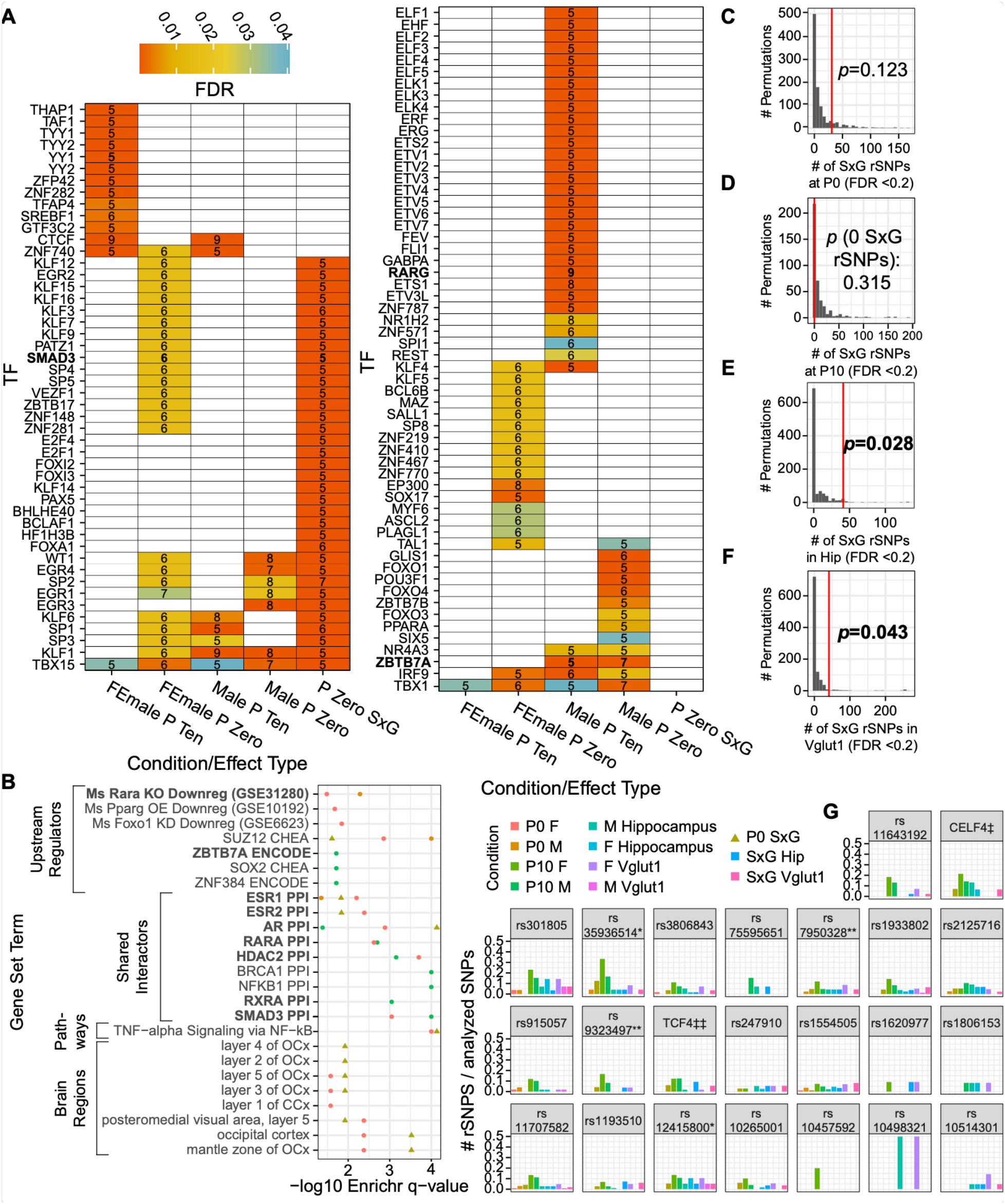
Regulatory architecture of rSNPs at P0 and P10, permutation analysis evaluating the number of detected SxG rSNPs, and the context-dependent landscape of MDD loci. (**A**) TFs with binding motif disruptions by ≥5 nominally significant (*p*_emp_ < 0.05) rSNPs or SxG rSNPs, enriched relative to nonfunctional SNPs. (**B**) Enrichr analysis findings for rSNP-enriched TFs in P0 and P10. Color key is shared with panel G. (**C-F)** Distribution of significant (FDR < 0.2) SxG rSNPs in 1,000 permutation analyses per condition (400 for P10); red lines: number of SxG rSNPs identified experimentally; overlaid *p*-values indicate the probability of observing as many SxG rSNPs by chance. (**G**) Each MDD locus, labeled by tag SNP, with bars representing the percentage of analyzed variants that were rSNPs or SxG rSNPs for each condition at *p*_emp_-derived FDR<0.2. Color key shared with panel B. *: locus from all-female MDD GWAS (*42*); **: near-genome-wide significant locus in males with MDD developing after age 50 (*41*); ‡: collapsed results from two loci near *CELF4* with tags less than 150kb apart; ‡‡: collapsed results from LD partners of five tag SNPs, comprising two GWAS significant tags and several weaker association peaks covering a span of ~5Mb around the gene *TCF4* (*6*).

Despite the absence of sex interactions in the P10 brain, we also found largely distinct sets of TFs in each sex at this age (only 4 shared) (**Fig. 6A**). Among the 42 P10 male TFs, 38 were male-specific, including *RARG*—again supporting our *in vitro* findings of retinoid-interactivity (*26*)— and several *KLF* members. (In contrast, *KLF*s were instead only enriched at P0 in rSNPs from *females*). The 15 female P10 TFs were predominantly core transcriptional machinery, including *YY1*/*2*, *GTF3C2, TAF1*, and in both sexes, *CTCF*.

We then looked at these TF sets as before to identify convergent regulators and functions among them (**Fig. 6B**; **Data S2D-E**). Of the few enriched annotations for male P0 TFs, we notably found that 4 corresponded to genes downregulated by *Rara* knockout, and 6 were putative regulatory targets of *PBX3*, which shows widespread subcortical and midbrain expression in E18.5 and P4 mouse brain (*74*). Female P0 TFs were enriched in Allen Atlas expression signatures for cortical layers 1, 3, and 5, and, as found in hippocampus, in PPI targets of *HDAC2* and *ESR1*. P10 male TFs showed the greatest extent of overlap (12/42) with gene-regulatory targets of *ZBTB7A* and were enriched for PPI targets of *HDAC2*, *RXRA* and *RARA*. The only enrichment found for P10 female TFs was via a modest (3/15) set of *FOXA1* regulatory targets.

#### Transcriptional-regulatory systems implicated in sex-differential rSNPs postnatally

Given the absence of FDR-corrected sex-genotype interactions at P10, we only analyzed P0 interaction rSNPs for TF motif perturbation enrichment. TFs enriched at interaction SNPs were comprised largely of *EGR*, *KLF*, and *SP* family members, as well as *PAX5* and neurodevelopmental factor *SMAD3* (**Fig. 6A**). Annotation of SxG TFs revealed a broader extent of hormonal roles in functional variation than observed in either sex alone at P0: we found SxG TFs were again enriched for PPI targets of *ESR1* (as had been P0 female TFs), but additionally enriched in PPI targets of *AR* and *ESR2*; the latter was otherwise absent from gene set analyses of TFs (**Fig. 6B**, **Data S2D-E**).

### Landscape of functional variation differs broadly across age, sex, and brain region/cell type

#### Sex-genotype interactions at MDD loci are overrepresented in hippocampus and its Vglut1^+^ neurons

Finally, we wanted to test the hypothesis that the number of SxG interactions from the assayed SNPs exceeded what was expected by chance. We randomly scrambled sex labels and repeated our analyses to define a distribution for the null expectation regarding number of SxG rSNPs at a *p*_emp_ FDR of 0.2. While our P0 SxG findings did not exceed chance expectations (**Fig. 6C**), our P10 results were consistent with a true absence of sex interactions (**Fig. 6D**). Most strikingly, the number of adult total hippocampal (**Fig. 6E**) and Vglut1^+^ (**Fig. 6F**) SxG interactions we identified were both significantly greater than would be anticipated.

#### Risk loci are characterized by multiple functional variants across age and cell type

Overall, we found that it was the norm for loci to contain multiple functional, context-dependent SNPs, in contrast to the concept of a singular “causal variant” driving a given GWAS association. Altogether, our analyses identified 280 rSNPs from 31 LD regions (28 depression-associated), with up to 13 rSNPs in a single locus found in a single condition (P10 female, tag SNP rs11707582 (*6*)) (**Fig. 6G**). In terms of age and cell type, we identified dozens of rSNPs with allele or SxG effects specific to one timepoint or tissue region: 26 P0-specific, 101 P10-specific, 34 total hippocampus-specific, and 55 specific to Vglut1^+^ cells of the hippocampus. Indeed, only 64 (~23%) of rSNPs are functional in more than one developmental or cell type context. Similarly, 92 rSNPs were only identified as functional in female conditions and 37 only in male (excluding SxG interactions)—or, excluding P10, 36 in female and 16 in male—while another 86 were only subject to SxG interactions. In other words, only 23% of rSNPs were sex-invariant in their *in vivo* activity.

## Discussion and Conclusion

We have directly measured sex-genotype interactions across MDD in the adult hippocampus and sexually differentiating brain, thus demonstrating the existence of a genetic component of sex differences in MDD and the regulatory architecture underlying these differences across space, time, and genome. We have uncovered functional differences between sexes at particular MDD-associated SNPs in the hippocampus, its excitatory neurons, and the brain during its sexual differentiation, expanding on observed genetic and clinical sex differences in MDD from general heritability to direct identification of sex-interacting variants.

Our data are supported by a wide variety of orthogonal datasets, including reporter assays and human brain epigenomic datasets beyond those used for variant prioritization. MDD-associated SNP rs1467013 was previously demonstrated to be functional in a classical luciferase reporter assay in three different cell lines (*75*). A prior *in vitro* MPRA identified rs301807, but not rs301806, as an rSNP (*26*), while both were identified as functional here. This highlights the importance of *in vivo* context for obtaining relevant insights about functional variation within GWAS loci—in this case, revealing two rSNPs in close (~2kb) proximity that likely influence expression of the same target (*RERE*).

Human datasets further support the translatability of our mouse approach in identifying regulatory SNPs and sex interactions. Four rSNPs identified here, rs7244124 (Vglut1^+^ SxG), rs76931017 (P10 female), rs827187 (P10 female), and rs4482931 (male hippocampus) were recently identified as chromatin accessibility QTLs in human midfetal neural progenitors and/or neurons (*76*), consistent with the allele-differential regulatory activity we observed. Intriguingly, an early attempt to identify sex-interacting GWAS loci for MDD (*77*) found suggestive significance for rs1345818, near *TMEM161B* and *MEF2C*, a locus which has since been sex-agnostically associated to MDD (*6*). Our assay did not include the reported sex-interacting GWAS tag, but we identified three variants in LD with rs1345818 that showed sex interactions, confirming that this risk locus indeed has sex-dependent regulatory activity: rs1814149 (hippocampal SxG, FDR<0.2, R^2^ with rs1345818=0.67), rs5869417 (hippocampal SxG, FDR<0.1, R^2^=0.21), and rs6452770 (Vglut1^+^ SxG, FDR<0.2, R^2^=0.69).

Downstream analyses aimed at identifying regulatory programs involved across rSNPs provided both support for the rSNP findings themselves, while also identifying novel candidate TFs underlying sex interactions at MDD loci. Our SxG-enriched TF sets were especially rich in sex hormone receptors and interactors, consistent with expectations for an *in vivo* assay detecting sex interactions, and indicating a role of sex hormone receptors in co-regulation of MDD risk variants. Our hippocampal analyses revealed male-specific roles for *AR*: male rSNPs were enriched for binding sequences of *ZMIZ1*, an *AR* co-activator, while an *AR* co-repressor, *ZBTB7A* (*78*), was identified as a shared upstream regulator of these TFs. Notably, *ZBTB7A* also regulates human non-coding RNA LINC00473 (*79*), which was recently been demonstrated to have sex-differentiated effects on depressive mouse behaviors when overexpressed in cortex (*80*). Our TF analyses of P0 SxG rSNPs also identified regulatory programs consistent with the critical period for sexualization of the brain. P0 SxG rSNPs were enriched for TFs interacting with *ESR1*, *ESR2*, and *AR*, consistent with the regulatory landscape necessary for accommodation of sex hormonal signals during the perinatal testosterone surge. Additionally, *PAX5* motif disruptions were unique to P0 SxG variants; interestingly the *PAX5* motif was recently shown to be enriched in promoters of sex-differentially expressed genes in adult brain (*21*). Our neurodevelopmental rSNP-enriched TFs likewise recapitulated aspects of recent preclinical studies of sex and depression: or example, P0 female rSNPs were enriched in motifs of endothelial marker *SOX17*, consistent with demonstration of sex-differential changes in mouse brain vascular permeability after stress (*81*).

Our assay has several limitations. Notably, our P0/P10 assays could not be compared directly to our adult findings to look for sex-by-age effects, as adult experiments were limited to the hippocampus while developmental assays were brain-wide. Unfortunately, it is impractical to region-specifically deliver AAV9 *in utero*, and hippocampal morphogenesis is postnatal. Likewise, episomal AAV delivery may not capture all regulatory information of a genomic delivery, though benchmark studies indicate a strong correlation (*82*). Finally, results might be influenced by use of either a different minimal promoter or longer fragments if nearby elements interact with the rSNPs. However, none of these limitations would be expected to create spurious sex effects, indicating the surprising and widespread SxG interactions are likely to be robust.

The presented *in vivo* MPRA approach indicates that critical biological and environmental factors involved in brain gene regulation and regulatory variation can be studied using a high-fidelity model of development, cell types, and biological signals. Our approach provides a framework for direct, functional study of psychiatric risk genetics and their interactions with biological and environmental factors that are imperfectly modeled *in vitro*, including cell type, sex, and brain development. This same approach could be readily used in the future to directly identify variants subject to genetic-environmental interactions with other key psychiatric risk factors, such as early life adversity and chronic stress.

## References and Notes

## Acknowledgments

Scott Lee for assistance with immunofluorescence of pilot hippocampal injections; Michael Vasek, Ph.D. and Lexi Harris for assistance with hippocampal dissections; Ernesto Gonzalez of the Washington University in St. Louis (WU) Hope Center Animal Surgery Core for performing hippocampal AAV9 delivery; EZH Switzerland Viral Vector Facility (VVF) facility for Sanger sequencing and AAV9 packaging of the MPRA library; McDonnell Genome Institute (sequencing), funded by National Institutes of Health grant UL1TR002345; WU Center for Genome Sciences and Systems Biology (sequencing); Barak Cohen, Ph.D., Tony Fisher, Ph.D., Tomás Lagunas, Jr., and Stephen Plassmeyer for insightful discussions and methodologic guidance; WU Center for Cellular Imaging (Axioscan microscope); Biorender (hippocampus experiment schematic); WU Epigenome Browser (*RSRC1* locus visualization); and Michael White, Ph.D. and Rachel Rahn, Ph.D. for reviewing the manuscript.

## Author contributions

Conceptualization: BM, JDD, DS

Methodology: BM, DS, JDD

Investigation: BM, DS

Visualization: BM, DS

Funding acquisition: BM, JDD

Project administration: BM, JDD

Supervision: BM, JDD

Writing – original draft: BM, DS, JDD

Writing – review & editing: BM, JDD, DS

## Data and materials availability

The annotation of library SNPs performed at the time of design are available at https://bitbucket.org/jdlabteam/n2a_atra_mdd_mpra_paper/src/master/Library%20Design%20Epigenomic%20Annotations/. Code is available at https://bitbucket.org/jdlabteam/paper-resources-mdd-in-vivo-mpras/src. Raw sequencing files and tabulated barcode counts per sample will be made available through GEO accession GSE186348 on publication. Outside datasets used for annotation are detailed in Supplementary Materials.

## Supplementary Materials (*83*–*115*)

Materials and Methods

Supplementary Text

Figs. S1 to S7

Tables S1 to S3

References (*##*–*##*)

Data S1 to S3

