## Supplementary material for "Sex significantly impacts the function of major depression-linked variants *in vivo*": Methods and Supplemental Text

**In this PDF:**

Materials and Methods
Supplementary Text
Figs. S1 to S7
Tables S2 to S3
Captions for Data S1 to S4

**Other Supplementary Materials for this manuscript include the following:**

**Table S1:** *Table S1-Sequencing Prep and QC Metrics.xlsx*
**Data S1:** *Data S1-Single Condition and Interaction P, Pemp, q-FDR, and signif T-F at FDR threshes, and single condition analyses betaA2 values.xlsx*
**Data S2:** *Data S2-Aggregated Significant Enrichr Results driven by 3plus enriched TFs for hippo and dev conditions.xlsx*
**Data S3:** *Data S3-Downsampled P10 n5 per sex LMM Results and SxG signif at FDR thresholds.xlsx*

### Materials and Methods

Animal Research Statement

All procedures involving animals were approved by the Institutional Animal Care and Use Committee at Washington University in St. Louis, MO.

Design and construction of minimal promoter-reporter-WPRE 3' UTR cassette.

A previously designed MPRA reporter consisting of a minimal *hsp68* promoter and dsRed-Express2 (*1*) was PCR amplified with a forward primer adding a 5' *MreI* cut site and a reverse primer with 3' overhang homologous to the 5' end of the WPRE 3'UTR element. The WPRE 3' UTR element was PCR amplified from a lentiviral plasmid encoding cyan fluorescent protein with the WPRE element (derived from plasmid FCIV) (*2*) with a forward primer containing a 5' overhang homologous to the 3' end of dsRed and a reverse primer adding a 3' *PacI* cut site. These PCR products were cut and purified from a 1% agarose gel and subjected to 10 cycles of PCR stitching (without primer) to allow overhangs to anneal and act as primers to create a contiguous sequence, followed by addition of hsp68-dsRed forward primer and WPRE reverse primer and 15 further PCR cycles to amplify the contiguous product. The PCR reaction was run out on a 1% agarose gel and the properly sized band (~1.5kb = ~1kb hsp-dsRed + ~500bp WPRE) was cut and purified. Later, Sanger sequencing (*described below*) confirmed proper assembly of the cassette.

An analogous version of this reporter was subsequently created using edited primer sequences to amplify the full cassette, replacing the 5' *MreI* site with a 5' *BsiWI* site and the 3' *PacI* site with an *AsiSI* site. This reporter was subsequently cut and cloned into single-oligo plasmids generated during the cloning optimization process. Single clones were grown in liquid culture and isolated plasmids were verified by Sanger sequencing to provide a clonal stock of reporter for the later full-scale MPRA library of psychiatric GWAS loci.

Design, construction, delivery, and RNA collection: cell-type-specific promoter proof-of-principle MPRA library.

*Proof-of-principle library barcoding and cloning*

Three promoters were PCR amplified with primers adding an *MluI* cut site to the 5' end, and *MreI* and *PacI* cut sites to the 3' end (for later insertion of the hsp-dsRed-WPRE cassette).

Promoters were **1)** a 2.2kb human *Gfap* promoter region (*3*) **2)** a 1.3kb mouse promoter region of the excitatory, neuron-specific gene *Camk2a*, amplified from plasmid pAAV.CamKII(1.3).eYFP.WPRE.hGH (Addgene plasmid # 105622, a gift from Karl Deisseroth to Washington University's Viral Vector Core); and **3)** a 521bp human promoter region of the constitutive, highly transcribed gene *Pgk2*, amplified from plasmid pRRLsinPGK-GFPppt (*2*). These PCR products were used as the template for a second PCR, in which a reverse primer homologous to the added *MreI and PacI* cut sites was used. The reverse primer also contained an overhang consisting of a 9bp region of N's, resulting in random sequences for use as barcodes, followed by a *SalI* cut site for insertion of the product into a plasmid backbone. For the minimal-promoter only condition, a single oligonucleotide consisting of all four cut sites, intervening bases to ensure cuttability, and a barcode (*MluI*-*MreI*-*PacI*-barcode-*SalI*) was ordered, and PCR amplified. For all promoter PCR products, barcode identities were determined by Sanger sequencing after insertion of PCR product into vector (see below).

Plasmid JD386 (*4*), originally encoding *mTdTomato* under the *Gfap* promoter, was digested at *MluI* and *SalI* sites (products #R3198 and #R3138, New England Biolabs, Ipswich, MA, USA) to remove the promoter and *mTdTomato*, leaving an intact human growth hormone (hGH) poly-A signal sequence just downstream of the *SalI* site, as well as intact ampicillin resistance and inverted terminal repeats necessary for AAV packaging. The promoter PCR product was digested with the same enzymes. The plasmid digest was treated with antarctic phosphatase (AP*)* (NEB #M0289) and gel purified to isolate the desired backbone fragment. The digested PCR products and gel-purified, AP-treated backbone digest fragment were ligated using T4 ligase (NEB #M0202) at 16°C for 18 hours with vector:insert molar ratios ranging from 1:3 to 1:5. Ligations were then directly transformed into DH5α chemically competent cells (NEB #C2987H), outgrown for 45 minutes in a 250rpm shaker at 37°C, then plated on LB agar with ampicillin and allowed to incubate for 16-18 hours at 37°C. Individual colonies were used to incubate 1mL wells of LB-ampicillin liquid media in 96-well deep-well plates and shaken for a further 16 hours at 37°C. The liquid cultures were then used to generate two identical 96-well plates of 25% glycerol culture stocks, one of which was sent for Sanger sequencing. Sequencing in this round of cloning used primers upstream of the *MluI* site and downstream of the *SalI* site, allowing verification of the promoter sequence and identification of barcodes. Up to 10 clones per promoter were selected based on having both expected sequencing results and unique barcodes. Equal volumes of each clone's glycerol stock were then used to inoculate a 10mL LB-ampicillin liquid culture of each promoter's "library 1" and grown at 37°C for 18 hours.

The "library 1" cultures were then mini-prepped (NucleoSpin Plasmid kit #740588.250 Macherey-Nagel, Düren, Germany) to isolate plasmids for insertion of the previously mentioned reporter cassette. Each library 1 plasmid pool, as well as the reporter cassette, was digested with *MreI* and *PacI* (Thermo Fisher #ER2021, Waltham, MA, USA; NEB #R0547). As before, the plasmid digest was treated with AP and gel purified. The digested, AP-treated, purified plasmid fragment and digested reporter cassette were combined at a vector:insert ratio of 1:5 and ligated using T4 ligase as above. Ligations were directly used for transformation, plating, and single-clone 96-well Sanger sequencing and glycerol stocking as described above. Sanger sequencing was performed with the same primers as for Library 1; this confirmed the presence of the inserted promoter (sequencing downstream from *MluI* site), as well as retention of the barcode and the presence of WPRE and the 3' end of dsRed insert (sequencing upstream from the *SalI* site). Clones were selected for generation of "library 2" based on present and intact promoter, minimal promoter, dsRed, and WPRE, and were only selected if the barcode identified was one of the barcodes from among the library 1 clones. This resulted in 5 to 9 eligible clones for each promoter condition. The clones' glycerol stock wells were again used as inoculum for a 10mL LB-ampicillin starter culture, used to inoculate 500mL LB-ampicillin cultures for maxiprep (Plasmid Maxi kit (#12163), Qiagen, Hilden, Germany). The four maxiprepped "library 2" plasmid pools (one pool per promoter) were submitted to the Washington University Viral Vectors Core for packaging into AAV9.

*Neonatal mouse transduction with cell-type promoter MPRA library(ies); subsequent TRAP and immunofluorescence (IF)*

One litter of SNAP25-RPL10a-eGFP TRAP mice (described in (*5*)), back-crossed for >10 generations to wild-type C57BL/6J mice, was genotyped at postnatal day 2 (P2). GFP-positive pups received intracranial injection of a mixture of all four viruses mixed at an equal titer; an aliquot of the mixture was set aside for later sequencing for DNA counts during MPRA analysis. GFP-negative pups (3) were injected with either the *hsp68*-only virus, *CAMK2A* promoter virus, or *GFAP* promoter virus to confirm cell-type specificity by immunofluorescence.

All animals were injected with 2µL bilaterally of virus or viral mix into three coordinate pairs: four anterior to bregma (two anteromedial, and two ~1.5mm posterolateral to the former), and two medially midway between bregma and lambda.

At P17, the three GFP-negative pups were perfused with 4% paraformaldehyde in PBS, brains dissected and dehydrated, and sliced in the coronal plane into 40µm floating sections stored in PBS with 0.01% sodium azide for immunofluorescence (further IF method details below). Tissue was stained with primary antibodies as follows: mouse anti-NeuN 1:500 (MAB377), goat anti-*GFAP* 1:250 (Abcam ab53554), and rabbit anti-dsRed 1:500 (Rockland 200-301-379). Secondary antibodies were used at 1:1000 donkey anti-mouse with Alexa Fluor 647, donkey anti-goat with Alexa Fluor 488, and donkey anti-rabbit with Alexa Fluor 546. Sections were then slide-mounted and imaged on a Zeiss LSM 700 (Zeiss, Germany) using a 40x oil objective.

At P27, the three surviving GFP-positive mice were sacrificed for individual biological replicates of TRAP (methods below), performed as previously described (*6*, *7*). Briefly, whole brain was separately homogenized from each mouse. 2.5% of the supernatant from the 20,000 *x g* spin of homogenate was collected as an "input" sample of RNA, i.e. RNA from all cell types in the tissue. Both input and TRAP RNA were QC'ed using an Agilent Tapestation's high-sensitivity RNA assay (Agilent, Santa Clara, CA). Two replicates were retained, with input RNA integrity number estimates (RINe) 7.8 and 8.2; TRAP RINe values were 6.8 and 7.4.

<u>Design, construction, delivery, and RNA collection: Neuropsychiatric GWAS MPRA library</u>

*Neuropsychiatric MPRA Library Design*

The Neuropsychiatric GWAS MPRA was designed as previously described (*8*). Briefly, tag variants were selected from neuropsychiatric GWAS studies, predominantly those of MDD or meta-analyzing MDD alongside additional neuropsychiatric disorders (*9*–*17*). Additional tag variants were selected from GWAS of anxiety disorders (*18*), attention deficit hyperactivity disorder (*19*), educational attainment/intelligence (*20*, *21*), and traits showing strong SNP coheritability with MDD (i.e., neuroticism (*22*, *23*), and mood instability (*24*)). As a negative control locus, we selected one tag related to anthropomorphic traits, rs1883640 (*25*). Loci were expanded to SNPs with LD $R^2$>0.1 and minor allele frequency >0.01 with the tag variant in their appropriate 1000 genomes phase 3 population (EUR for all studies except two loci from the Han Chinese CONVERGE GWAS of MDD) using LDLink (*26*). SNPs were manually selected for inclusion in the MPRA library on the basis of overlap with both human brain eQTL (*27–32*) *and* at least one additional human brain epigenomic annotation (*33–38*). 11 SNPs were removed due to introduction of a nonsynonymous coding change. Two loci instead included all SNPs in LD $R^2$ >0.65 and SNPs with a RegulomeDb (*39*) score of ≥ 4 for LD $0.1 < R^2 < 0.65$. For loci especially sparse in overlaps with the screening annotations, a single overlap of any sort was considered adequate. We note that while this approach is not wholly empirical, recent machine learning-based approaches have identified that regulatory variation is better predicted by considering annotations only in the variant's local genomic region (as we did manually here).

Oligonucleotides (oligos) were designed for the 1453 selected SNPs, containing up to 126bp of human genomic sequence (hg19) centered on the variant. For small insertion/deletion variants, the larger allele was used to define the 126bp sequence (i.e., the smaller allele was 126-(allele length difference) bp). Each allele of each variant was assigned 10 randomly generated 10bp barcodes from a filtered set ≥ 2 Hamming distances apart, comprised of 25-75% GC, and filtered for runs of >3 of any one base. Promoter-only ("basal") oligonucleotides instead included a 126bp filler in a region of the oligonucleotide cut out during cloning (below), paired to 110 of these barcodes. The deeper barcoding of the basal oligonucleotide allows for a well-powered, within-sample normalization factor to be calculated (see analysis methods below). The remaining oligonucleotide sequence, 200bp in total, consisted of restriction sites for cloning and primer sites for oligo amplification from the single-stranded oligo pool (**Fig. S1**).

A 200bp oligonucleotide pool, comprised of 29,280 unique sequences, was ordered from Twist Biosciences (San Francisco, CA). The oligonucleotides were flanked on the outside by a forward priming sequence, 5' GAGGGAAATCGTGACGCGTG 3' (forward primer same sequence), and reverse priming sequence of 5' GTCGACCAGGTCATCACTATTG 3' (reverse primer 5' CAATAGTGATGACCTGGTCGAC 3'). The internal portions of these sites are MluI (followed by a G, to prevent creation of other used restriction sites resulting from design of adjacent genomic sequence) and SalI cut sites, respectively. Proceeding from the 5' end, the remaining space comprised the ≤ 126bp allelic sequence of interest, followed by BsiWI, PmeI, and AsiSI sites end-to-end, and the barcode (**Fig. S1**).

*Cloning of the MPRA library*

Oligonucleotides were amplified by PCR for 12 cycles in 6, 50µL reactions, using Phusion High-Fidelity Polymerase 2x Mastermix (New England Biosciences) each with 500nM final primer concentrations and 10ng oligonucleotide pool as template. Reactions were thermocycled as follows: 98°C, 3 minutes → 12 cycles of 98°C/15s, 65°C/30s, 72°C/15s → 72°C/5min → 4°C. Products were loaded into an unstained, 2% tris-acetate-EDTA (TAE) agarose gel run at 90V for 105 minutes. The gel was post-stained with 15µL 10,000x SyBr gold (Invitrogen) with 150mL of TAE in a small pyrex dish, gently rotated for 20 minutes, and the 200bp product band cut for gel purification using the Nucleospin Gel and PCR Clean-Up kit (Takara Biosciences, Kusatsu, Shiga, Japan). 12ng of product was purified for cloning.

The purified PCR product wascut in a 100µL double digest reaction containing 1.5µL MluI-HF (NEB), 1.5µL SalI-HF (NEB) and supplied buffer at 37°C / 1hr followed by an 80°C/20 min heat kill. The AAV-compatible plasmid backbone (JD386, see above) was digested separately in four 100µL reactions, each containing 2µL MluI-HF, 2µL SalI-HF, and 2µL BamHI-HF (to resolve the target product on subsequent gel), supplied buffer, and 2µg plasmid, incubated at 37°C / 1hr and heat killed at 80°C / 10min. Subsequently, 4µL H2O, 4µL Antarctic Phosphatase, and 12µL of supplied buffer (NEB) were added to each reaction; the reactions were then divided into 60µL aliquots for further incubation at 37°C / 1hr. The vector was then run on a 0.8% agarose gel with GelGreen dye (Biotium, Fremont CA), the target 3.4kb band cut and purified using the Nucleospin kit per manufacturer directions. The oligo digest reaction was used directly for ligation with the purified vector at a molar ratio of vector:insert 6:1. Nineteen 20µL ligation reactions were aliquoted from a single master mix containing 11.9ng digested oligo, 1.51µg digested, phosphatased, and purified vector, 10µL Enzymatics T4 ligase and 40µL supplied buffer (Qiagen). Ligations incubated at 16°C / 14hr, then heat killed at 80°C / 15min. Purification of the ligation product was performed using MyOne Silane magnetic beads (Thermo Fisher) from which the supplied buffer was removed and replaced with 3x the total ligation volume of Qiagen Buffer QG. The ligation reactions were recombined into a single 1.5mL microcentrifuge tube with the buffer QG•silane bead mixture, and incubated end-over-end at room temperature for 75 minutes. Beads were then washed on a magnet stand by removing supernatant and adding 80% ethanol thrice (without disrupting magnetized beads). The beads then air dried for 10 minutes and product was eluted into 196µL of water, with 192µL taken for transformation into *E. coli*.

For transformation, 6µL (~18 ng) of ligation product was added to each of 27 tubes of 50µL of DH5alpha chemical competent *E. coli* (NEB), which were transformed per manufacturer instructions. Three transformations were simultaneously performed using DH5alpha electrocompetent *E. coli* (NEB), each consisting of 8µL ligation product (~24ng) and 100µL cells in a 1mm cuvette per manufacturer instructions. All transformants were grown out for 45 minutes at 250 rpm and 37°C in their PCR tubes (chemical transformation) or after transfer into culture tubes with 2mL pre-warmed SOC (NEB; electrochemical transformation). The outgrowths were pooled (19mL total), added to a flask of 31mL luria broth (LB) with carbenicillin for a maxiprep starter which incubated with 250rpm, 37°C shaking for 6 hours. The full starter volume was then added to 450mL LB with carbenicillin and cultured with 250rpm shaking at 37°C for 10 hours. The plasmid was maxiprepped using GenElute HP Plasmid Maxiprep Kit (Sigma Aldrich, St. Louis MO). Low-depth next-generation sequencing was performed on the maxiprep with amplicons generated over the space spanning the MluI and SalI sites to confirm the presence of the library.

The Sanger-verified *hsp68*-dsRed-WPRE cassette (described above) was cut back out of its backbone in four 30µL digest reactions, each containing 2µg of the clonal miniprep plasmid, 1µL BsiWI-HF (NEB), 1µL AsiSI (NEB), supplied buffers, heat killed at 80°C/10min, then gel purified from a GelGreen-stained 0.8% agarose gel using the Nucleospin kit. The maxiprepped "first plasmid library" was digested in 20µL reactions of 215ng each, using 1µL AsiSI for 37°C/1hr, followed by direct addition of 1µL BsiWI-HF, 1µL 10x CutSmart buffer, and 8µL water, and digestion at 37°C for a further hour. 1µL rSAP phosphatase (NEB) and 9µL H2O was then added to the reaction and incubated at 37°C / 1hr, 80°C / 20min, and brought to 4°C. Forty 20µL ligation reactions were prepared as described above, each containing ~50ng DNA in 1:6 molar ratio of the cut vector and reporter cassette. The ligations were thermocyled 25°C/30s, 16°C/30s 20 hours (~800 cycles in practice) followed by 80°C/20min heat kill and 4°C incubation. Ligations were cleaned up with Silane beads as described above.

The "second plasmid library" was transformed into 25 tubes of DH5a chemically competent cells; 20 reaactions were 200µL cells, 19µL (~100ng) ligation; 2 were ~150µL cells and 5µL (~25ng) ligation; 2 were 50µL cells and 5µL ligation, and one was 50µL cells and

2.5μL (~12.5ng) ligation, all of which were transformed per manufacturer instructions. Cells were outgrown for 50 minutes at 37°C/250rpm. Outgrowth was pooled and used to inoculate a starter and subsequent maxiprep culture as described above. Sequencing (in the same manner as performed for later experimental analyses) was then performed to verify barcode coverage of the plasmid library. The resulting plasmid was sent to EZH Zurich Viral Vector Facility, which verified its sequence by Sanger sequencing and packaged the library in AAV9.

*Hippocampal Stereotaxic Delivery*

Hippocampal stereotaxic injections were performed in a counter-balanced fashion (alternating order male/female, with the order switched each day) over a three-day period (n=4 per day). Mice were anesthetized with continuous isoflurane up to 5% until unreactive to toe pinch, then maintained at 2-3%. The skin of the head was retracted to expose the surface of the skull. A 0.485 μM hamilton syringe was stereotactically guided into place and used with an automatic pump to deliver 2μL of AAV9 at a rate of 0.2μL per minute to each of four positions (8μL AAV9 total per animal; deeper coordinate first): A/P -2.0 mm, M/L -1.5mm and 1.5mm, and depth (from dura) of -1.75mm first, followed by partial withdrawal of the syringe and an additional delivery at -1.25mm depth. The needle was allowed to dwell for 10 minutes after each injection was completed. Mice were then given intraperitoneal buprenorphine 0.1mg/kg and cranial subcutaneous lidocaine 25mg/kg for post-operative analgesia and recovered in the home cage under a heat lamp with cagemates. Mice were checked for ambulation and absence of distress at 1, 24, 48, and 72-hours after return to consciousness. Mice then continued to live in the home cage with same-sex cagemates (same cagemates as prior to surgery) with the same *ad libitum* access to food and water and the 12:12 light/dark cycle they had been reared under.

*In-Utero AAV injections and brain collection/lysis:*

CD1 IGS timed-pregnant female mice were purchased and delivered (Charles River Laboratories). Pregnant dams were allowed to house overnight after delivery to reduce stress and ensure optimal success during in-utero injections. Pregnant mice at 15 days of gestation were placed under isoflurane anesthesia and a midline laparotomy was done to expose the uterus. 1uL of viral MPRA-AAV was mixed with 0.025% Fast Green FCF (Sigma, St. Louis MO) and administered through a 10uL-Drummond glass micro dispenser pipette (Drummond Scientific,

Broomall, PA) into pups through the uterine wall. Injections were targeted towards the anterior horn of the lateral ventricles as previously described in IUE and AAV injection studies (*40*, *41*). For each dam, 2-4 pups were left non-injected due to position of pup in-utero or impaired development compared to other pups. On P0 and P10, mice were sacrificed and screened for fluorescence under a dual fluorescent protein flashlight (NightSea, Lexington MA). P10 mice brains were weighed post-removal to assess long-term changes in brain weight due to viral injection. Pups were decapitated and brains dissected out with removal of the cerebellum, and the remaining brain placed in a microcentrifuge tube on dry ice, then stored at -80°C overnight to aid homogenization. The following day, the brains were removed from -80°C storage for addition of 500µL (P0) or 1mL (P10) Trizol reagent, homogenized thoroughly using a battery-powered hand pestle, and returned to -80°C until all samples were collected for that age group for single-batch RNA purification (*below*).

Translating Ribosome Affinity Purification (TRAP)

For immunoprecipitation (IP) of GFP-containing ribosomes, 60µL per sample of streptavidin MyOne T1 beads were resuspended in 17µL of 1µg/µL/sample protein L (reconstituted in 1x PBS), 36µg/µL/sample anti-eGFP 19C8 and 36µg/µL/sample anti-eGFP 19F7 (both available through Sloan-Kettering's antibody & bioresource center https://www.mskcc.org/research/ski/core-facilities/monoclonal-antibody-core-facility), brought up to 200µL times the number of immunoprecipitations to be run with 1x PBS. The beads in this mixture were incubated with end-over-end mixing at 4°C for two hours to bind antibodies to the magnetic beads. Beads were then separated on a magnet stand and washed by resuspension and remagnetization 5 times using 0.1% bovine serum albumin (BSA) in 1x PBS. Beads were then washed three more times using wash buffer (above), then resuspended to a final volume of 105µL/IP and kept on ice until needed.

On the day of tissue collection, DTT, RNAse inhibitors, and cycloheximide were added to stock TRAP buffers to the specified concentrations detailed below. Mice were deeply anesthetized with isoflurane, rapidly decapitated, and the brain removed and dissected for TRAP. Each brain was blunt dissected to remove anterior and posterior-most portions, then placed in a pre-chilled dish on ice containing 15-25mL of modified TRAP-compatible buffer consisting of 1x phosphate-buffered saline (PBS), 0.1 mg/mL cycloheximide (to halt translation for ribosome capture), and 1/2500 vol/vol each of rRNAsin (Promega, Madison, WI USA) and SUPERase•in (Thermo Fisher, Pittsburgh, PA USA). Hippocampus was dissected out bilaterally in a dish of buffer on ice, then both hippocampi from a single animal homogenized to constitute a sample. Samples were sequentially dissected and homogenized.

TRAP was then performed as previously described with slight modifications. A wall-powered drill, run at full speed, was fitted with a Teflon pestle was used to homogenize each sample in 1mL pre-chilled homogenization buffer (10mM HEPES pH 7.4, 150 mM KCl, 10mM $MgCl_2$, 0.5 mM diothiothreitol (DTT), 0.1 mg/mL cycloheximide, and 1/1000 vol/vol each of rRNAsin and Superasin, and Roche EDTA-free protease inhibitor cocktail (dissolved in homogenization buffer without DTT, cycloheximide, or RNAse inhibitors) to a final concentration of 1x). Homogenates were spun down at 2,000 x *g* for 10 minutes at 4°C. 825µL of supernatant was collected and combined with 100µL each of 10% NP40 in water and 300mM 1,2-diheptanoyl-sn-glycero-3-phosphocholine (DHPC; Avanti Polar Lipids, Alabaster, AL USA) and incubated on ice for 30 minutes. This solution was then spun at 20,000 x *g* for 15 minutes at 4°C. For the input (or here, "total hippocampus") RNA fraction, 50µL of supernatant was collected and added to 200µL of wash buffer (1% vol/vol NP40, 10mM HEPES pH 7.4, 150 mM KCl, 10mM $MgCl_2$, 0.5 mM diothiothreitol (DTT), 0.1 mg/mL cycloheximide, and 1/1000 vol/vol each of rRNAsin and Superasin, and Roche EDTA-free protease inhibitor cocktail (dissolved in homogenization buffer without DTT, cycloheximide, or RNAse inhibitors)) and 750µL Trizol LS (Thermo Fisher), and stored at -80°C until all TRAP and input samples had been collected.

975µL of the same supernatant was taken for ribosome capture by IP and added to an aliquot of 100µL of the resuspended, antibody-coupled beads. The mixture then incubated end-over-end for 5.3-5.7 hours at 4°C. After incubation, beads were separated on a magnet stand, supernatant removed, and resuspended in 1mL high-salt wash buffer (10mM HEPES pH 7.4, 350 mM KCl, 10mM $MgCl_2$, 0.5 mM diothiothreitol (DTT), 0.1 mg/mL cycloheximide, and 1/1000 vol/vol each of rRNAsin and Superasin, and Roche EDTA-free protease inhibitor cocktail (dissolved in homogenization buffer without DTT, cycloheximide, or RNAse inhibitors)) before remagnetizing. This supernatant was removed, the beads suspended in a second wash of high-salt buffer, and transferred to a new microcentrifuge tube (to avoid nonspecific RNA stuck on tube walls from releasing by later addition of Trizol LS). Beads were then magnetized and washed twice more with the high-salt wash buffer. Beads were then resuspended in 250µL of the *wash* buffer (i.e., the 150mM KCl buffer described), and 750µL Trizol LS was added. These samples too were stored at -80°C until all samples were collected.

RNA purification (all experiments) and brain DNA isolation (P0/P10)

RNA purification was using the Zymo Clean and Concentrator-5 (Zymo, CA USA), with simultaneous processing of all samples for each condition (i.e., all hippocampal input and TRAP samples in one batch; all P0 samples in one batch; all P10 samples in one batch, see **Table S1**). The samples were removed from -80°C, allowed to come to room temperature for 5-10 minutes, followed by addition of 20% volume of chloroform. Tubes were shaken vigorously by hand for 30 seconds and allowed to stand for 7 minutes at room temperature. Subsequently, tubes were spun at 12,000 x *g* for 20 minutes at 4°C. For hippocampal samples, 500µL of supernatant was collected; otherwise, 175µL was collected. The remaining phase-separated Trizol-chloroform sample was returned to -80°C (see DNA collection below).

For uniform RNA handling, the Zymo kit instructions were followed, but prepared one mastermix adequate for all samples being purified so as to avoid variability in volumes of buffer/ethanol added to each. This mastermix consisted of 2 supernatant volumes of Zymo RNA Binding Buffer and 3 supernatant volumes of 100% ethanol per sample. 5 supernatant volumes of this mix was added to each Trizol-chloroform supernatant, mixed thoroughly by pipetting, and applied to the kit columns, spun through at 12,000 x *g* at room temperature until columns were loaded. Manufacturer instructions were then followed for the remainder of cleanup. RNA quality was assessed using Agilent High-Sensitivity RNA Tapestation assay. All samples in all experiments used for sequencing preparation, including the proof-of-principle pilot, had an RNA integrity number (RIN) of, at minimum, 6.

For isolation of DNA from P0 and P10 brain to verify the presence of MPRA barcodes at the DNA level, phase-separated Trizol-choloroform mixtures were brought to room temperature and used for DNA isolation according to manufacturer instructions.

Immunofluorescence:

Brains not isolated for RNA analysis were removed and fixed in paraformaldehyde in PBS (4%) followed by serial sucrose in PBS solutions (15%, 30%). Following post-fixation, brains were embedded in OCT (Sakura, Torrence CA) and sectioned at 30µm (hippocampus) or 35 µm (P0, P10) using a Leica CM1950 cryostat (Buffalo Grove, IL) and processed for immunofluorescence as slide-mounted sections for P0 and free-floating sections for P10 pups and adult hippocampus.

To characterize adult hippocampal AAV9 delivery, 30µM coronal sections were cut 21 days after delivery, incubated in 1x PBS with 5% normal donkey serum and 1:1000 chicken anti-GFP (to identify TRAP-positive cells), 1:500 rabbit anti-RFP ((1:500, Rockland, 600-401-379; to identify AAV-transduced cells), and 1:500 goat anti-GFAP (1:500, Abcam ab53554) to visualize potential astrocytosis around the viral injection sites. Slices incubated in primary antibodies overnight at room temperature on a horizontal mixer, were rinsed three times in 3x PBS, followed by 45 minutes of incubation in 1x PBS with 5% normal donkey serum and 1:1000 each of Alexa Fluor 488 donkey-anti-chicken, Alexa Fluor 568 donkey anti-rabbit, and Alexa Fluor 647 anti-goat. Sections were rinsed again with 1x PBS, then incubated for 5 minutes in PBS with 1:20,000 DAPI for nuclear fluorescence, rinsed once more with 1x PBS, then mounted onto slides with application of Prolong Gold, followed by nail polishing of cover slips into place. Slides were stored in a foil-covered box at 4°C until imaged on the Axioscan.Z1 slide scanner (ZEISS, Germany) at 10x resolution.

For P0 Primary antibodies included anti-RFP (as above), anti-GFAP (as above), and anti-NeuN (1:250, MAB377). Fluorescently conjugated secondary antibodies (AlexaFluor 488, 568, and 647) were obtained, and nuclei were labeled with a DAPI counterstain. “No primary” controls were done for both sets of time points to indicate and ensure staining was specific to primary antibody targets. Multi-channel imaging was performed at 20X using an AxioScan.Z1 slide scanner to assess both independent region and whole-brain viral transduction.

Image editing was performed using ImageJ software and only included re-scaling of resolution, brightness/contrast adjustments, and cropping.

<u>RNA-seq library prep (proof-of-principle and Neuropsychiatric GWAS MPRA libraries)</u>

RNA samples were treated with the Turbo DNA-Free kit (Ambion #AM1907, Austin, TX, USA) to remove extant DNA using the manufacturer’s instructions for high-concentration DNA (2µL of enzyme, followed by 20% volume of Inactivation Reagent to remove the DNAse). Sequencing libraries were prepared from RNA by performing a variation on the methods of *e.g.*

(*1*, *42*, *43*) by using a reporter-specific primer, targeting the polyA signal sequence just 3' to the barcode sequence (*44*), during first-strand reverse transcription with Superscript III Reverse Transcriptase (Invitrogen 18080044, Carlsbad, CA, USA). Double-stranded cDNA was then synthesized and amplified by PCR using Phusion HF (NEB #M0531) using the same reverse primer and a forward primer in the WPRE of the 3'UTR. These primers added unique cut sites allowing subsequent Illumina adapter ligation. To prevent sequencer errors due to homogenous sequence at the start of read 1 (3' end), these adapters were a mix of four different lengths to stagger the first base of the 3' end read. Digestion, clean-up, ligation, clean-up, and final PCR with primers with partial homology to the adapter ends were used to add the remaining Illumina sequences and sample indices. The full details of each sample processing step, including bead-based size selection, were as described (*8*). Sample input mass, number of PCR cycles for the two PCR steps (single-strand cDNA to double stranded DNA and index PCR), and read depths are described in **Table S1**. Sequencing of proof-of-principle samples was performed on a MiSeq instrument (Illumina, San Diego, CA); all other experiments were sequenced on an Illumina NovaSeq 6000 instrument. NovaSeq 6000 (Illumina).

qPCR verification of cell-type marker changes in TRAP

20µL reverse transcriptase reactions were prepared using Quanta Biosciences qScript with supplied 5x buffer (containing both random hexamer and poly-T primers), containing 20ng of sample RNA. Reverse transcriptase reactions were incubated at 25°C/5minutes, 42°C/30minutes, 85°C/5minutes, then kept on ice. RT reactions were diluted with 140µL water (final volume 160µL) for use as qPCR template. 10µL qPCR reactions (technical triplicates per sample•gene) were prepared using Sybr Green 2x Mastermix (Thermo Fisher), containing 4µL cDNA, 0.5µL each primer, and 5µL of qPCR mastermix. qPCR thermocycling ran for 40 cycles at 95°C/15s, 63°C/30s per cycle, followed by a melt curve on a Quantstudio 6 instrument. Before analysis, data were quality checked by a) examining melt curve product heights (all were a singular, consistent peak per gene over 80°C, corresponding to a true amplicon as opposed to primer dimers) and b) identification of outlier wells based on a cycles to threshold of detection (CT) value ≥1 cycle different from other sample•gene technical replicates. One row of technical replicates corresponding to a total hippocampal sample were removed due to large CT discrepancies relative to the other two technical replicate sets; 6 other singular wells were excluded on the same basis of outlier status, and 1 well was excluded for failure to amplify any product. All analyzed sample•gene wells contained at least two technical replicates in strong agreement (CT values within 0.5 of one another).

To assess enrichment in TRAP relative to total hippocampal RNA, the technical replicate mean CT value for the internal control gene, ß-actin (*Actb)* was subtracted from the CT value of each other gene. qPCR primer sequences are in **Table S2**.

MPRA Sequencing analysis

*Proof-of-principle experiment*

Barcodes were counted from read 1 sequences, allowing up to 3 mismatches in the 20bp upstream of the barcode and 0 mismatches within the barcode sequence itself. The number of reads mapping to each barcode were totaled and normalized to counts per million (CPM) with normalization for sequencing library size (in number of reads mapped) using EdgeR (*45*, *46*). Expression for a given barcode was then calculated as the ratio of (CPM RNA / CPM viral DNA) for each RNA sample (TRAP and input RNA from each brain). Expression values were normalized to the within-sample mean expression of the minimal promoter (*hsp68*) alone by taking expression (BC in sample) / expression (mean(*hsp68* BCs)) ). By normalizing within sample (i.e., input or TRAP), expression is thus normalized to general minimal promoter activity for that sample . Significance was calculated by performing repeated-measures ANOVA / linear mixed modeling. Each barcode group's expression was implemented as a repeated measure and modeled as a dependent variable of RNA sample type and of a random variable for source tissue, thus: *expression ~ RNA.fraction + (1|mouse)* (*47*). For plotting and interpretation of barcode enrichment/depletion between biologically paired input and TRAP RNA samples, log2 fold-change in expression was calculated by subtracting log2(normalized expression in Input) from log2(normalized expression in TRAP) for each barcode.

*Sequencing analysis: All other experiments.*

Barcodes were counted from read 1 sequences, allowing mismatches neither within the barcode sequence nor the 6bp upstream/8bp downstream of flanking sequence. CPM were calculated using the total number of barcode-mapping reads prior to several filtering steps; for samples with multiple sequencing runs of data (DNA technical replicates and a subset of P0 RNA samples), a single CPM value per barcode per sample was first calculated by obtaining the mean CPM across the sequencing runs. 1) Barcodes with a DNA count under a specified threshold (187 or approximate CPM equivalent) are excluded from the counts table from the DNA *and* RNA samples. 2) For sequences with 4 or fewer remaining DNA barcodes represented across all samples after this step, all other barcodes for the sequence are excluded from all of the samples in the table to avoid analysis of sequences with inadequate barcoding depth. 3) Counts of RNA barcodes are then removed on a per-barcode-per-sample basis if they fall below a separate minimum read threshold (75 counts or approximate CPM equivalent), set below the DNA threshold to allow for detection of repressive effects. 4) Preliminary expression values for barcodes (log2 (RNA barcode CPM / DNA barcode CPM)) are calculated for each replicate and collapsed across barcodes into a mean for each Regulatory Element (RE) within sample. Single barcodes with outlier expression values (≥ 2 standard deviations) apart from other barcodes for that sample are dropped only from that sample. (The expression values are not written out to a results table at this time). 5) Penultimately, all barcodes are dropped from individual samples if 4 or fewer barcodes remain for a given sequence in that sample, such that all samples analyzed for a given sequence have at least 4 barcodes represented in each sample. 6) A final check is made to ensure that each barcode remaining is represented in at least 50% of samples, and those represented in fewer samples are removed, followed by a second check that all samples have ≥4 barcode expression values remaining.

Expression for a given barcode was then calculated as the log2 ratio of (CPM RNA / CPM viral DNA) for each RNA sample. Prior to linear modeling, within-sample barcode expression values were normalized by subtracting the within-sample mean expression of the set of barcodes paired to the minimal promoter (*hsp68*) alone (<=110 barcodes). By normalizing within sample, expression is thus normalized to minimal promoter activity among the cell types and proportions comprising the RNA sample—while this notably does not change inter-sample correlations, it *does* alter the Euclidean distance between samples in hierarchical clustering (namely, samples with outlying barcode wise expression before normalization cluster back in with the other samples after this transformation). Shapiro tests for normality were performed on each condition-wide set of barcode expression values for each SNP (i.e., 4-10 barcodes * N samples * 2 alleles); all Shapiro tests were >0.05 for all analyzed SNPs in all eight conditions.

Linear mixed modeling was then applied within single sexes for each condition to analyze allelic effects alone, and with data from both sexes pooled to test for allele-by-sex interactions. Each barcode group's expression was implemented as a repeated measure and modeled as expression as a dependent variable of allele (and for interaction models, of sex and sex-by-allele), thus: *expression ~ allele + sex + allele*sex + (1|barcode)*. We note that the random intercept values determined for barcodes were consistent even when running the model using different samples, indicating we were detecting and removing biologically invariant effects of the barcode sequences on RNA levels (**Figs. S2-S4**). Empirical null test statistics were calculated as previously described (*8*); in brief, we applied the respective experimental model to 50,000 comparisons between two "alleles" each comprised of 6 randomly selected barcodes from among the 110 corresponding to the *hsp68* minimal promoter alone, thus controlling for the noise inherent to the assay and sample set. These values were then used in the qvalue package (*48*) to determine empirical p values, and corresponding q-values and FDR significance were generated from the empirical test statistics and p-values (also using the qvalue package).

Analysis of functional SNPs for enrichment in TF motifs

To assess TF binding sites potentially disrupted by functional variants, we first generated sets of positive (functional) and negative (non-functional) variants for the analyses. For single-sex, single age/tissue analyses, we defined functional SNPs as those with an uncorrected $P_{emp}<0.05$, and the remainder of the measured SNPs as non-functional. We also performed two comparisons for each age/tissue to identify TFs enriched at sex-genotype interaction SNPs. One comparison assessed interaction SNPs at $P_{emp}<0.05$ to "non-functional" SNPs ($P_{emp} > 0.05$ for both allele and sex-allele interaction) to maximize the size of the negative set used in enrichment analysis. We additionally compared functional interaction SNPs to those with only a significant main effect allele effect, so as to identify TFs potentially involved in sex-divergent variant effects. The enrichment procedure required a negative set of greater size than the positive set (a random set of negatives equal to the size of positives was drawn each iteration, see below); to meet this condition, we allowed for more lenient definition of allele-only effects from the LMM. The thresholds and number of SNPs in the positive and negative sets are shown in **Data S3A**.

We performed motif perturbation analyses for all SNPs designed into the MPRA utilizing two tools: 1) the R package motifbreakR(*49*) and its built-in database of motif position-weight matrices (PWMs) from multiple public repositories, and 2) RSAT var-tools (*50*) with each of 3 motif databases; its own 2017 database, comprised of clustered motifs (some without TFs assigned) based on similarities across multiple motifs for multiple TFs(*51*), JASPAR 2020 TF-specific motifs (*52*), and cisBP 2017's human database (*53*), which consists of both specific TF motifs and more general motifs not assigned to any one TF. Both tools are designed to identify PWM matches overlapping input SNPs in dbSNP (version 151 in hg38 for motifbreakR) or Ensembl (hg37, for RSAT) for which at least one of the SNP alleles results in a genomic sequence significantly matching a given motif sequence. We used the default significance cutoff of $p < 10^{-4}$ for calling motif matches in all analyses and identified changes in motif match score using the tools' default algorithms. MotifbreakR considers a weighted sum based on the position weights of each base in the motif sequence and considers these for the two alleles of the query SNP; the magnitude of these differences is used to classify motif perturbations as "strong" or "weak". For motifbreakR, we performed separate enrichment analyses only considering those changes classified as strong, and regardless of the algorithm's classification. RSAT performs a similar analysis, identifying the strongest position-weighted p-value match to a motif for each allele of a SNP within a sequence of user-defined length (here, we used 122bp flanks to approximate the 110-126bp sequences assayed in the MPRA) under a first-order (dinucleotide) background frequency model, and reporting the best match p-value for each allele to each motif where at least one allele exceeds the defined cutoff for the best match value.

Frequencies at the level of TF (which can include several motifs) were considered as the number of SNPs matched to a given TF, regardless of the number or identity of motifs to which that SNP matched. Null distributions of frequency were determined by 50,000 random selections of *n* SNPs of motif perturbations identified in the negative SNP set, where *n* was the number of positive set SNPs analyzed. Due to incomplete compatibility of hg37 and UK Biobank rsIDs with hg38/dbSNP 151, only 1277 SNPs were actually analyzed by motifbreakR; 1452 were analyzed by RSAT; the number of positive SNPs analyzed, and negative SNPs drawn in permutations were based on the number of SNPs actually analyzed in each respective tool. The p-value of frequency was then calculated from the empirical percentile of the positive SNP frequency count vs the distribution of frequencies in the negative sets, and these were corrected using standard FDR correction within each individual analysis, with resulting significant enrichments considered as FDR<0.05. We additionally logged whether each TF/motif was depleted in the positive SNP set relative to the permuted negative sets.

Finally, some results from the RSAT analyses using the RSAT 2017 and cisBP human 2017 databases corresponded to motif sequences not ascribed to particular TFs (by nature of those databases). We separated these enrichment results from those for which a TF was explicitly listed in motifbreakR or RSAT. To predict corresponding transcription factors for the undefined motifs, we utilized the MEME-suite tool TomTom (*54*) to predict significant matches (using Euclidean distance) of the database motif to TF-specific motifs across 4 databases: JASPAR CORE Vertebrates 2018 (non-redundant) (*55*), Jolma 2013 (*56*), Mouse Uniprobe (*57*), and HOCOMOCO v11 (human and mouse motifs) (*58*). Corresponding TFs were assigned for all TomTom matches at $p < 10^{-4}$; for cisBP/RSAT motifs where no TomTom match achieved this p-value, the single-lowest match p-value under 0.01 was retained.

Gene Set Enrichment Analyses of Motif-enriched TFs

From the above analyses, we generated lists of unique TFs identified across the 3 RSAT and motifbreakR analyses for each condition (9 hippocampal TF sets total--Vglut1 sex-genotype interaction, hippocampus sex-genotype interaction, both interaction types combined, and allelic rSNPs from each sex in each of tissue fraction and in both tissue fractions for each sex). For the neurodevelopmental conditions, we generated one TF set per age•sex, and one TF set for sex-by-allele effect variants from the P0 condition.

To narrow the hippocampal TF sets down to those most likely present (expressed) and thus able to exert regulatory activity in the adult hippocampus, we collected the publicly available GTEX v8 transcripts per million (TPM) expression dataset and subsetted to hippocampal samples (see *data availability* below), averaging the genewise TPM values across all samples (for interaction TF filtering) or against single-sex sample sets (for single-sex allele-effect TF filtering). From those TFs significantly enriched (FDR<0.05) from the analyses above—including those identified by matching nonspecific database motifs to putative TFs with TomTom—we filtered down to those with ≥ 3 average TPM in the respective GTEX hippocampal sample set. We did not perform any expression filtering of the P0 or P10 TF sets as comparable whole brain datasets do not exist.

We then utilized Enrichr to identify gene sets across ontologies, pathways, and drug perturbations where the TFs were enriched as a set (only considering enrichments of reported q-value < 0.01 and driven by ≥ 3 input genes if either the input gene or the result gene was a retinoid receptor or sex hormone receptor). **Data S3C and S3E** provide links to the enrichr results for each of the 9 GTEX-expressed and rSNP-enriched TF sets from hippocampus and for the 5 neurodevelopmental TF sets.

Permutation tests of sex-by-allele interactions

In order to determine the null expectation for the rate of significant sex-genotype interactions, we performed 1,000 iterations of each sex-by-genotype linear mixed model, wherein the sample labels were randomly shuffled by sex. The same linear model, including the 50,000-iteration empirical p-value calculation step as described above, was performed for each permutation to ensure that the empirical p-values were of the same granularity as the experimental analyses. The 20% FDR cutoff for the empirical p-values were determined for each iteration, and the number of SNPs significant for a sex-allele interaction at this threshold were recorded for each iteration. The end result was a vector of 1,000 numbers of FDR 20% "sex-genotype" significant rSNPs from the permutations, constituting a null distribution of the number of interaction effects for a given experiment. That was than compared to the actual number of interaction SNPs found with the true labels.

### Supplementary Text

Further information from *in utero* AAV delivery

We additionally confirmed at collection that brain weights were not significantly different between dsRed-positive and negative pups at P10 (**Table S3**). MPRA sequencing of P0 and P10 samples revealed a much greater frequency of outlier samples in terms of early quality metrics (lower number of barcodes recovered, low pairwise count correlations). We urge future users of this technique to consider collecting twice the number of samples desired for analysis; fortunately, wild-type pregnancies allow for this type of throughput.

Annotation datasets and other outside datasets

•GTEX v8 TPM https://storage.googleapis.com/gtex_analysis_v8/rna_seq_data/GTEx_Analysis_2017-06-05_v8_RNASeQCv1.1.9_gene_tpm.gct.gz ; de-identified metadata used to identify hippocampal samples https://www.ebi.ac.uk/arrayexpress/files/E-MTAB-5214/E-MTAB-5214.sdrf.txt
•Hi-C contacts for dopaminergic neurons and cortical neurons: https://github.com/thewonlab/H-MAGMA/blob/master/Input_Files/Midbrain_DA.genes.annot and https://github.com/thewonlab/H-MAGMA/blob/master/Input_Files/Cortical_Neuron.genes.annot
•Brain Hi-C contact matrices from Jung 2019: ftp://ftp_3div:3div@ftp.kobic.re.kr
•Song 2019 *in vitro* neural cell type and fetal primary astrocyte Hi-C: Corresponding paper's Supplementary Table 2
•Song 2020 fetal radial glia, intermediate progenitor cell, excitatory neuron, and inhibitory neuron chromatin contacts: files "iN.MAPS.peaks.txt", "IPC.MAPS.peaks.txt", "RG.MAPS.peaks.txt", and "eN.MAPS.peaks.txt" thru BDbag linked at https://assets.nemoarchive.org/dat-uioqy8b
•Su 2021 Hi-C data for neural tissues: ebi.ac.uk with accession E-MTAB-9159
•Fetal cortical plate and germinal zone Hi-C contacts from Won, 2016: corresponding paper's supplementary tables S22 and S23.

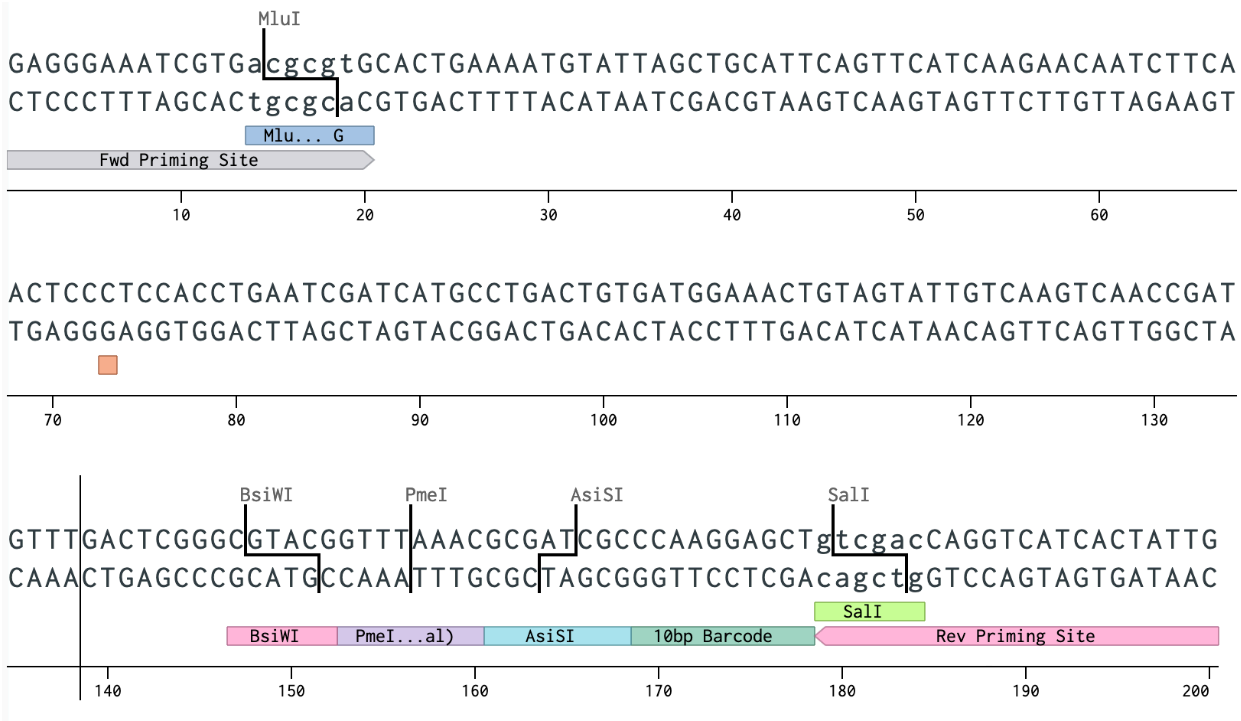


**Fig. S1. Template MPRA oligonucleotide.**

Example MPRA oligonucleotide illustrating the features described including priming and cloning sites. The red box underlines the variant position at the center of the 126bp human genomic sequence tile. The PmeI site serves as a failsafe in the design such that, should a high fraction of plasmid not take on reporter constructs during cloning, digestion linearizes the reporter-negative plasmids and retransformation of the digested DNA results in isolation of reporter-positive plasmids. (This was not necessary for this library). Illustration captured from sequence design tools bundled with digital lab notebook, Benchling (www.benchling.com).

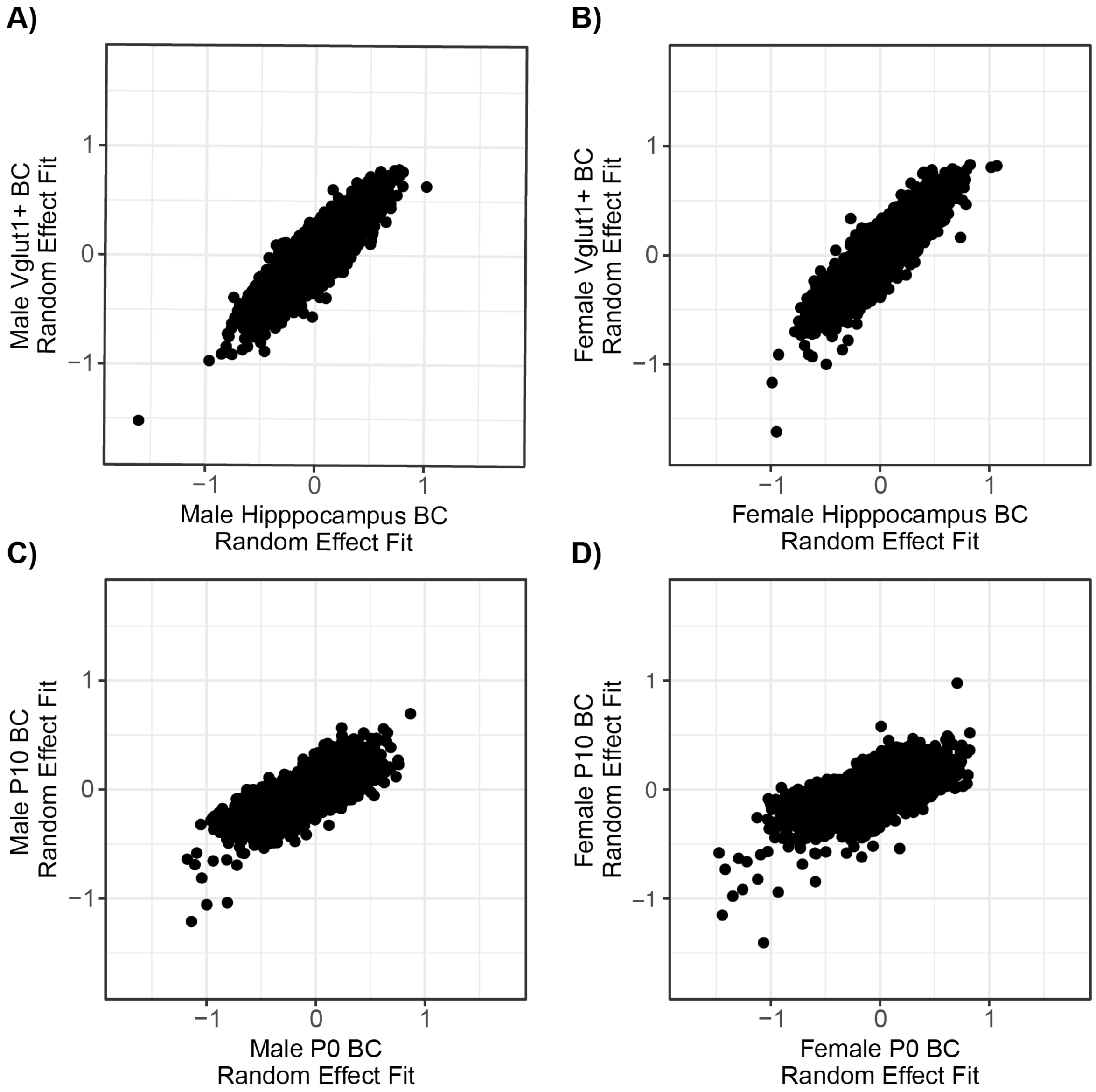


**Fig. S2. Barcode random effect coefficients are consistent within sex across ages/cell types.** **A)** Random effect coefficients from separate LMMs used to analyze male total hippocampus and male Vglut1$^{+}$ MPRA data. **B)** *Ibid.* for female. **C)** Random effect coefficients from separate LMMs used to analye male P0 and male P10 MPRA data. **D)** *Ibid.* for female.

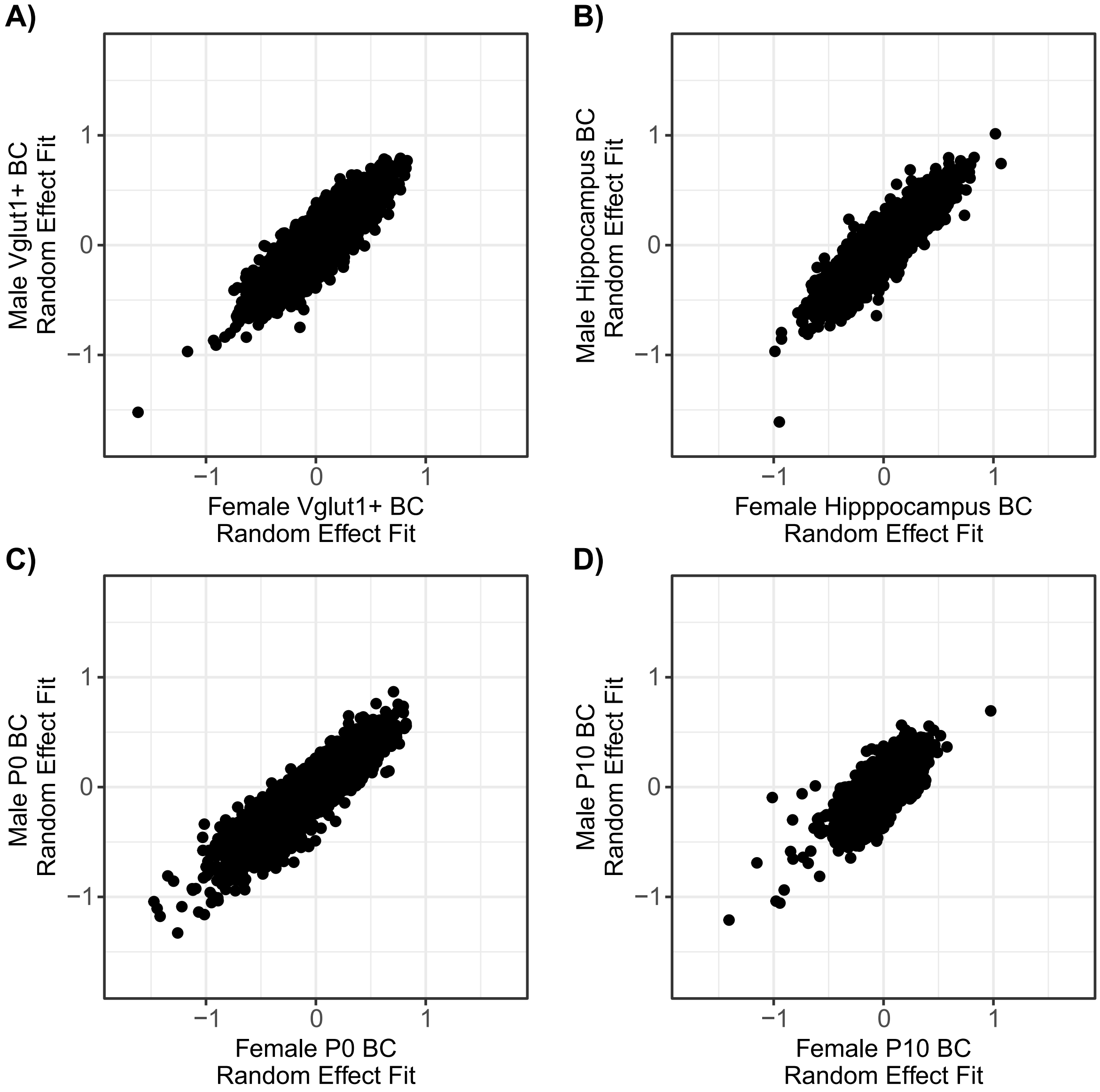


**Fig. S3. Barcode random effect coefficients are consistent between sexes.**
**A)** Vglut1$^{+}$ BC random effect coefficients from female vs. male. **B)** *Ibid.* for total hippocampus. **C)** *Ibid.* for P0. **D)** *Ibid.* for P10.

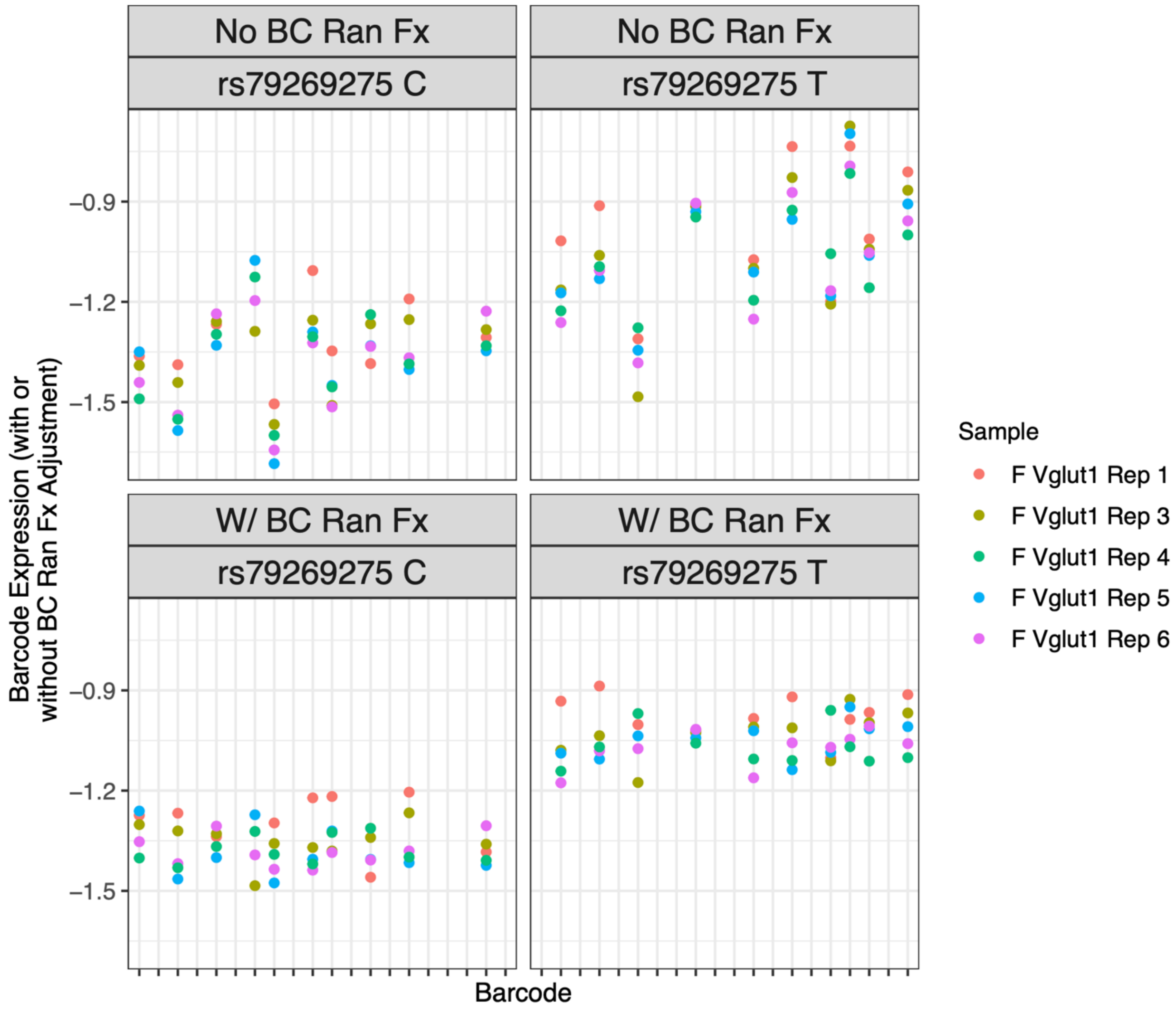


**Fig. S4. Example of barcode random effect fitting on expression values.**
Each X axis position is a barcode, with its expression level before or after adjustment for random effects shown by color for each replicate among female Vglut1$^{+}$ samples.

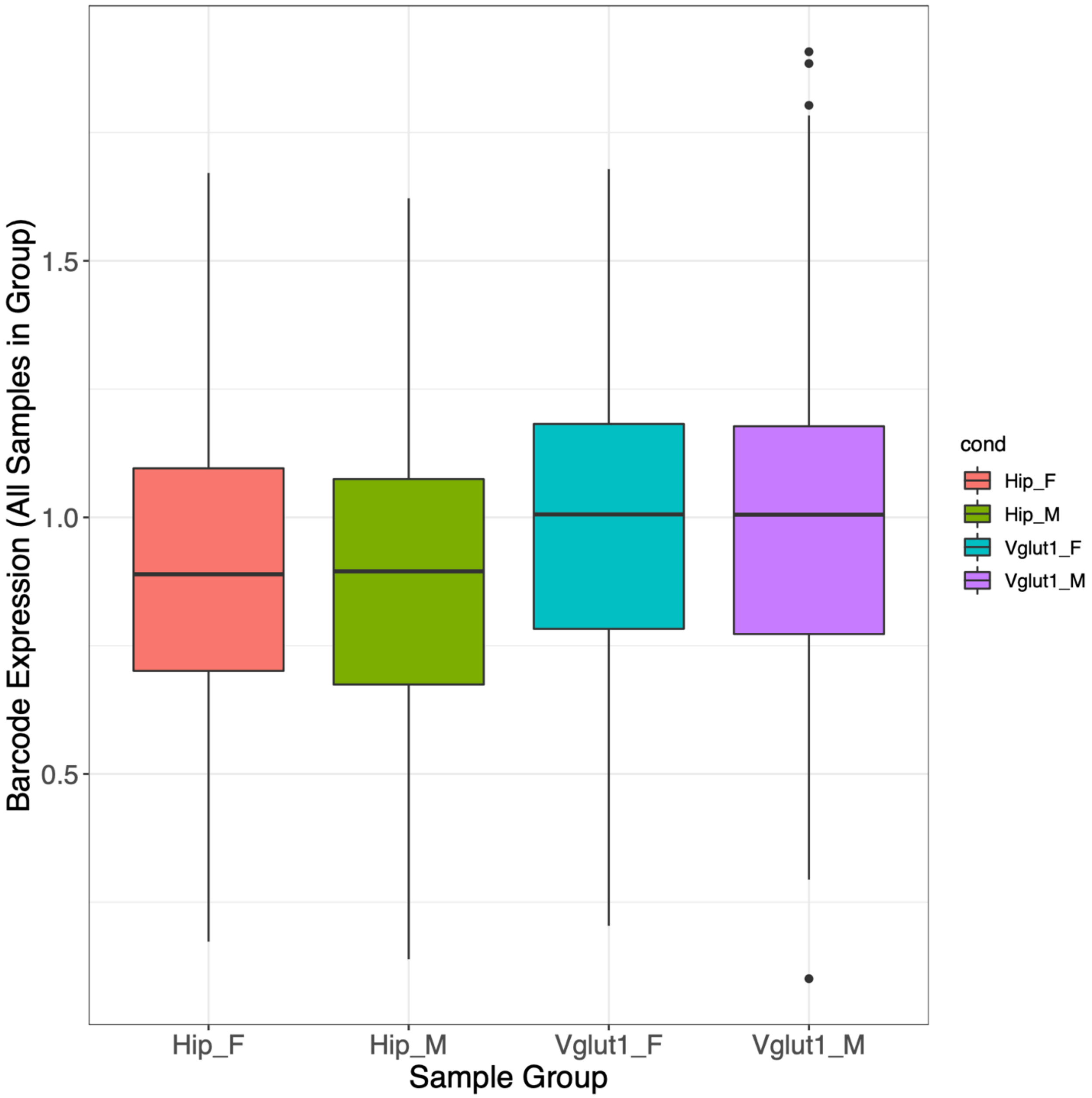


**Fig. S5. Basal (minimal promoter alone) barcode expression values do not vary by sex in total hippocampus or *Vglut1*$^{+}$ TRAP.**

P>0.5 for both sex comparisons using student's *t*-test; ≥ 99 BC expression values shown per sample type.

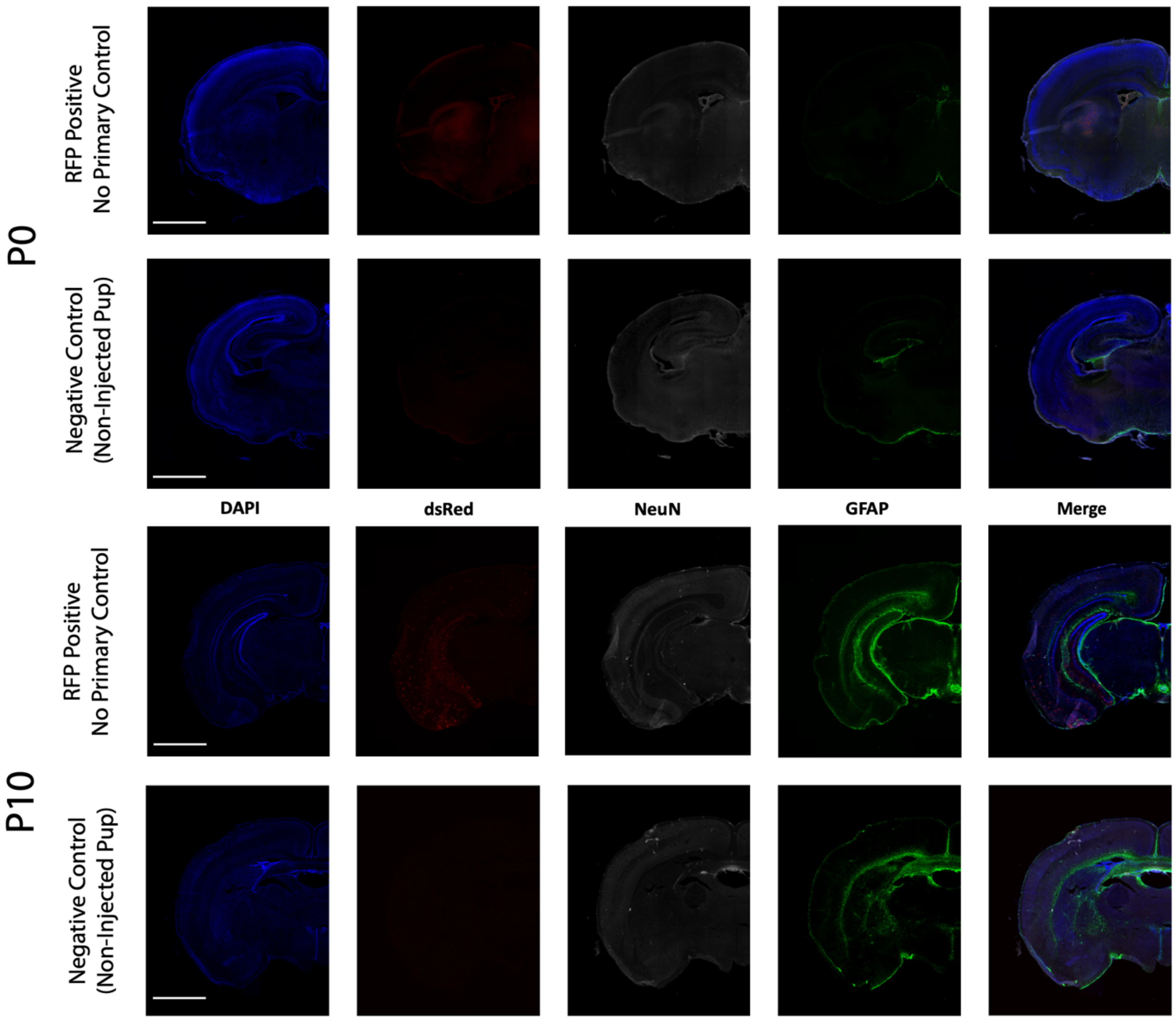


**Fig. S6. IF negative controls for P0 and P10 AAV9 delivery.**

**A)** An RFP-positive P0 brain without primary antibodies applied. **B)** An RFP-negative P0 brain *with* primary antibody staining (i.e., demonstrating that the dsRed primary antibody is selectively marking the AAV-delivered reporter per the main figure IFs). **C)** An RFP-positive P10 brain without primary antibodies applied. **D)** An RFP-negative P10 *with* primary antibody staining.

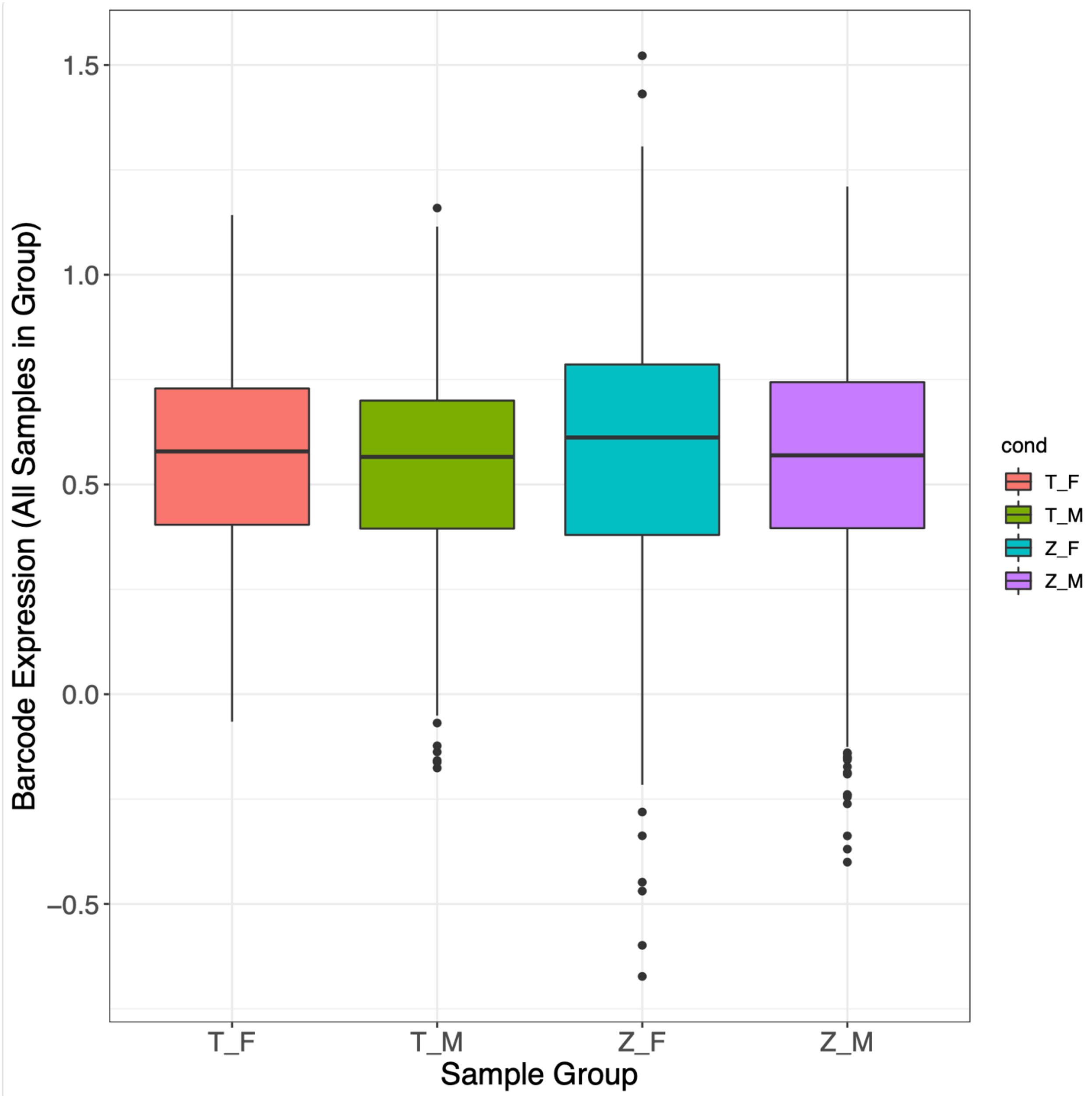


**Fig. S7. Basal (minimal promoter alone) barcode expression values do not vary by sex in P0 or P10 whole brain.**

P0 sex comparison p=0.14 (student's *t*-test); P10 sex comparison p=0.11 (student's *t*-test).

**Table S1. dsRed-positive and -negative (i.e., injected and uninjected) P10 littermate brain weights after *in utero* AAV-MPRA delivery.**

Table describing sequencing preparation parameters: starting mass of RNA or DNA, number of PCR cycles for each amplification step, and number of resulting samples for sequencing. Sequencing outcomes provided are the number of retained (QC-passing) samples for analysis, and read depth for RNA and DNA samples in each sequencing group.

**Table S2. qPCR primers for *Vglut1* TRAP validation (R=reverse, F=forward).**

| Target and priming direction | Primer sequence (5' -> 3') |
|---|---|
| Actb R | CAATAGTGATGACCTGGCCGT |
| Actb F | AGAGGGAAATCGTGCGTGAC |
| Snap25 F | CAACTGGAACGCATTGAGGAA |
| Snap25 R | GGCCACTACTCCATCCTGATTAT |
| Gfap Fw | AACCGCATCACCATTCCT |
| Gfap R | CGCATCTCCACAGTCTTTACC |
| Gria1 F | CAAGTTTTCCCGTTGACACATC |
| Gria1 R | CGGCTGTATCCAAGACTCTCTG |
| P2ry12 F | ATGGATATGCCTGGTGTCAACA |
| P2ry12 R | AGCAATGGGAAGAGAACCTGG |

**Table S3. dsRed + and - (injected or uninjected) P10 brain weights after *in utero* AAV.**

| Weight (g) | dsRed | Sex |
|---|---|---|
| 0.31 | + | M |
| 0.3259 | + | M |
| 0.3203 | + | M |
| 0.2816 | - | M |
| 0.2916 | + | M |
| 0.3679 | + | F |
| 0.3193 | + | F |
| 0.2973 | - | F |
| 0.3263 | - | F |
| 0.2824 | + | F |
| 0.3259 | - | F |
| 0.3261 | + | F |
| 0.2621 | + | F |
| 0.3254 | + | F |
| 0.3794 | + | M |
| 0.335 | + | M |
| 0.3406 | + | M |
| 0.3612 | + | M |

All negative pups are from litters that underwent *in utero* AAV delivery.

**Data S1. LMM reported p-values, empirical p-values ($p_{emp}$), derived FDR (q) values from each p-value type, and whether a SNP was significant for allelic or SxG effects at 5 $p_{emp}$ FDR thresholds; LMM allele (log2 fold-change) coefficients from each singly analyzed condition.**

Sheets, lettered A-U, are separated by analyses and significance/betas. Sheet A) lists LMM reported allele effect (single-condition analyses) or allele-sex interaction (interaction analyses) p-values, empirical p-values, and q-values (FDR) for $P_{emp}$. Significance data is also provided for all 8 single-condition analyses and the 4 SxG analyses in separate tables, with true/false status for 6 FDR cutoffs (5 $p_{emp}$-derived and FDR 0.05 for the LMM-reported p-value). Beta values are also provided for each of the 8 single-condition analyses in adjacent sheets. **All beta values are the effect of the listed 'A2' allele relative to the listed 'A1' allele** (i.e., negative ß signifies lower expression under the A2 allele compared to the A1 allele); these allele assignments are consistent across all analyses.

**Data S2. Motif enrichment SNP set significance thresholds, Enrichr analysis results involving ≥3 of the rSNP-enriched TFs, and links to the full Enrichr results in the web tool.**

**A)** Significance thresholds used for designating positive and negative SNP sets for each motif enrichment analysis. **B)** Results of the examined Enrichr gene sets (TF PPIs, upstream regulators, etc) tested for enrichment in each of the 9 discussed TF sets from adult TRAP-MPRA analyses. **C)** Links to the Enrichr web tool pages with results from the TF analyses (including the tool's built-in gene set analyses, the vast majority of which are undiscussed nor in data S2B or S2D. **D)** Results for the examined Enrichr gene sets tested for enrichment in each of the 5 discussed TF sets from developmental MPRA analyses. **E)** Links to the Enrichr web tool results for the developmental MPRA TF sets.

**Data S3. Results table from SxG LMM of the downsampled P10 cohort.**
